# Direct Optical Quantification of Chain Collapse, Reduced Dielectric, and Water Release Driving Protein Phase Separation

**DOI:** 10.1101/2025.11.04.686660

**Authors:** Ethan A. Perets, Jacob A. Spies, Justin H. Cheong, Lixue Shi, DeeAnn K. Asamoto, Alex S. Holehouse, Judy E. Kim, Wei Min, Jens Neu, Elsa C. Y. Yan

## Abstract

Biomolecular condensates represent unique microenvironments that organize intracellular biology and promote biochemical reactions. However, the biomolecular interactions driving condensate phase separation are often weak, transient, and heterogeneous. Investigating the structural biology and chemical properties of condensate interiors has therefore proven experimentally challenging, often requiring the use of perturbative probes. To overcome this challenge, we combine label-free optical scattering and vibrational spectroscopy approaches spanning ultraviolet, visible, mid-infrared, and terahertz wavelengths with deep-learning-based ensemble prediction of intrinsically disordered protein conformations. Our experimental and computational results reveal that the intrinsically disordered N-terminal domain of the RNA Deadbox helicase 4 (DDX4) deviates from random coil behavior and undergoes chain collapse that correlates with phase separation, which leads to lower dielectric and reduced water content inside condensates. Our data support a model of DDX4 phase separation whereby chain collapse, reduced dielectric, and water release enhance the strength of multivalent protein-protein interactions within condensates, driving condensate growth and phase separation through positive feedback. Our study addresses the critical driving forces of biomolecular phase separation across a range of length scales, providing quantitative insights into protein-protein/protein-solvent interactions and the chemical properties of condensate interiors.

## INTRODUCTION

Over the last decade, there has been growing appreciation that biomolecular condensates play crucial roles in the organization of intracellular biology.^1-6^ Biomolecular condensates are non-stoichiometric assemblies that concentrate specific biomolecules while excluding others. Condensates are often described as viscoelastic liquid-like assemblies, meaning that condensates flow like liquids but also possess rheological properties more commonly associated with elastic solids.^7^ Condensate liquid-like properties arise from multivalent interactions between condensate components, including cation-π interactions, π-π stacking, hydrogen bonding, electrostatics, and dipole-dipole interactions.^1,8-10^ Although these molecular interactions have been well-characterized in dilute conditions, how the internal environment of a condensate affects these interactions remains challenging to understand. Moreover, condensate interiors and interfaces are chemically distinct from their surrounding aqueous environment and from one another.^11-14^ Probing the internal molecular structure and chemistry of condensates in a way that does not destroy that environment remains a significant and outstanding challenge in the field.

The protein-protein and protein-solvent interactions that underlie condensate formation are generally weak, transient, heterogeneous, and therefore often difficult to probe using routine biochemical methods.^15^ Methods to probe these interactions often rely on the introduction of labels, including non-native fluorescent protein tags^16^ which can modulate the interactions that underlie assembly.^17,18^ Thus, label-free methods for probing liquid-like condensates are crucial to understand the native dynamic multivalent interactions associated with phase separation. Among such label-free methods, optical scattering and vibrational spectroscopy approaches are non-perturbative probes of biomolecular structures, dynamics, chemical microenvironments, interactions, and functions. In principle, such label-free optical methods could also be used to reveal the physical and chemical properties within condensates. Moreover, these approaches can interrogate protein-water interactions within the condensate interior, which remain understudied.^19-23^

Here, we investigated homotypic condensates formed by the intrinsically disordered N-terminal domain (NTD) consisting of the first 245 amino acids of the RNA Deadbox helicase 4 (DDX4^[1-245]^). Prior work established the DDX4 NTD as a model system for investigating the molecular driving forces that underlie condensate formation.^24-27^ We combine light scattering, electronic absorption, and vibrational spectroscopies, including Raman, infrared, and terahertz spectroscopies, spanning the electromagnetic spectrum from the ultraviolet (UV), visible (vis), mid-infrared (mid-IR), and terahertz (THz) regimes (**Figure 1**). Supported by computational analyses of protein conformational ensembles, UV-resonance Raman spectroscopy and confocal Raman microscopy reveal that DDX4^[1-245]^ remains disordered across dilute and condensed phases, but that the protein samples conformations that deviate from random coil-like behavior. Dynamic light scattering reveals the existence of a previously unreported DDX4^[1-245]^ collapsed state that correlates strongly with phase separation. Terahertz spectroscopy enables direct measurement of the reduced dielectric constant in the DDX4^[1-245]^ condensed phase. Finally, stimulated Raman imaging quantifies water release from DDX4^[1-245]^ condensates. Our results lead us to a model of DDX4^[1-245]^ phase separation whereby enthalpy-driven interactions promote protein chain collapse, leading to reduction of the local dielectric constant and water release that strengthens multivalent protein-protein interactions, likely creating positive feedback that can induce condensate formation and growth.

**Figure 1.**
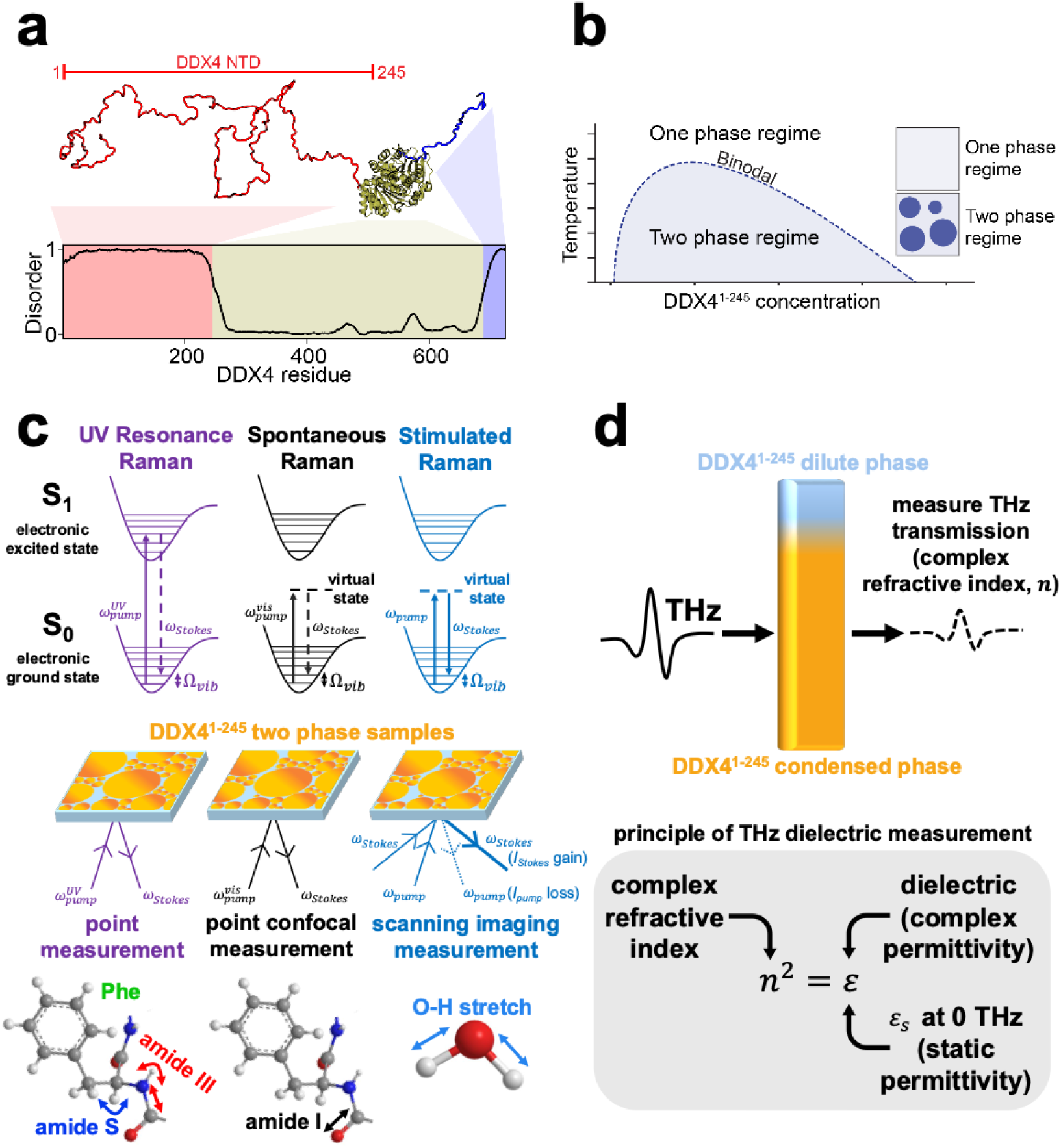
Vibrational spectroscopies to probe liquid-liquid phase separation of DDX4^[1-245]^. (a) Representative computational structural conformation and disorder of full-length DDX4, including the DDX4^[1-245]^ N-terminal domain (NTD) studied here. (b) An idealized phase diagram of DDX4^[1-245]^ phase separation. (c) Simplified Jablonski diagrams and observables for UV-resonance Raman, spontaneous confocal Raman, and stimulated Raman imaging. UV-resonance Raman point measurements detect DDX4^[1-245]^ phenylalanine (Phe) protein sidechain vibrations and determine the structure of DDX4^[1-245]^ by probing the amide S (protein backbone C_α_-H bending) and amide III vibrations (coupled protein backbone C-N stretching and N-H bending). Spontaneous Raman confocal imaging determines the structure of DDX4^[1-245]^ inside individual condensates by probing the protein amide I vibration (mainly protein backbone C=O stretching). Stimulated Raman imaging quantifies the water content inside individual DDX4^[1-245]^ condensates and at the condensate surface by probing the water O-H stretching vibration. (d) Terahertz (THz) spectroscopy measures the frequency-dependent complex refractive index (*n*) of the sample to extract the complex static permittivity (*i*.*e*., dielectric constant at zero frequency, ***ε***_***S***_) inside the DDX4^[1-245]^ condensed phase.

## RESULTS

### DDX4^[1-245]^ is intrinsically disordered but exhibits spectroscopic signatures that deviate from those of a random coil

Prior work has established that the DDX4 NTD forms homotypic biomolecular condensates, yielding DDX4 NTD molecules in the dilute and condensed phases (**Figure 1b**). Using nuclear magnetic resonance (NMR) spectroscopy, Nott *et al*. found that DDX4^[1-236]^ is an intrinsically disordered protein that remains disordered in the condensed and dilute phases after homotypic phase separation *in vitro*.^25^ Moreover, Brady *et al*. revealed that DDX4^[1-236]^ backbone dynamics in both the condensed and dilute phases undergo fast local reconfiguration.^26^ These prior works largely excluded that the DDX4 NTD possesses stable secondary or tertiary structure in the condensed or dilute phases. However, it is not known whether the DDX4 NTD behaves as a self-avoiding random coil (*i*.*e*., a chain with excluded volume but no meaningful attractive or repulsive interactions)^28^ or possesses sequence-encoded local and long-range interactions.

To investigate the conformational properties of DDX4^[1-245]^ in the dilute and condensed phases, we applied UV-resonance Raman spectroscopy and confocal Raman microscopy (**Figures 2a-c**). In UV-resonance Raman spectroscopy, the excitation wavelength is resonant with electronic transitions in the protein, making this technique a sensitive probe of aromatic protein sidechains and the amide backbone. In particular, the amide III mode, which is the coupled C-N stretching and N-H bending along the protein backbone, can be used to determine protein secondary structures.

**Figure 2.**
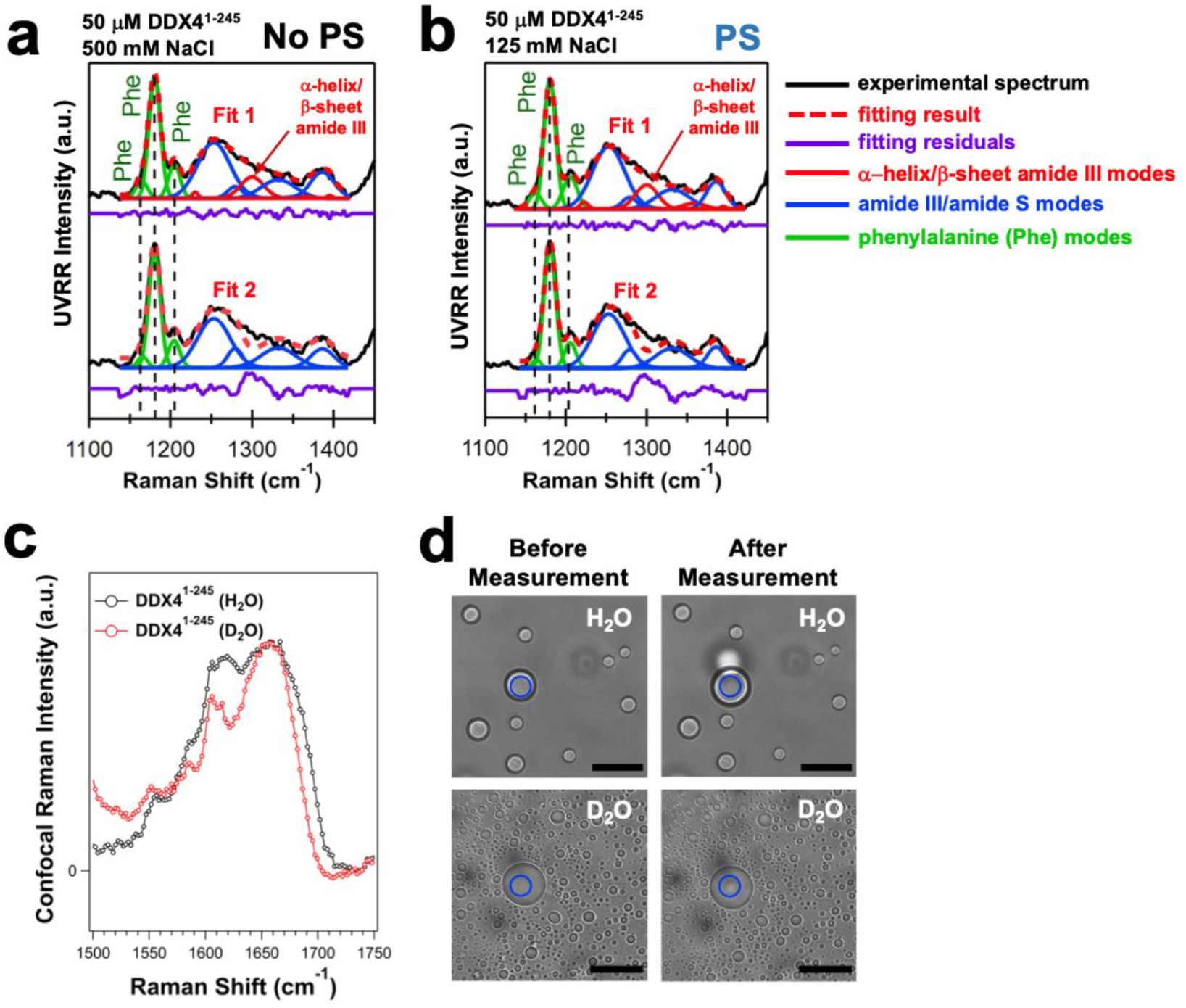
DDX4^[1-245]^ is an intrinsically disordered protein with non-random-coil structure. (a,b) UV-resonance Raman (UVRR) spectra (λ_ex_ = 215 nm, 5 mins. total acquisition, 25 °C) of (a) one-phase solution of DDX4^[1-245]^ (No PS) or (b) two-phase solution of DDX4^[1-245]^ condensates (PS). Signal from tryptophan was removed via subtraction of the UVRR spectrum of the model compound (L-)Trp. Black curves are experimental spectra, and Gaussian decompositions are shown. The red dashed curves are the sum of individual Gaussian vibrational contributions from phenylalanine (Phe) modes (solid green curves), amide III modes corresponding to α-helix/β-sheet structure (solid red curves), and additional amide S (1386 cm^-1^) and amide III modes (solid blue curves). The two data sets shown in each panel are the same experimental data fit either (top) with or (bottom) without modeling the α-helix/β-sheet-associated amide III vibrational response. The corresponding residuals from each fit are shown below each spectrum in purple. (c) Confocal spontaneous Raman spectra (λ_ex_ = 532 nm, 25 °C) of DDX4^[1-245]^ condensates prepared in (black) H_2_O and (red) D_2_O measured in the amide I vibrational region. Each spectrum is an average of 10 acquisitions (1 min. per acquisition). (d) Brightfield images of DDX4^[1-245]^ condensates, with regions measured in (c) marked by the blue circles. Samples are shown before and after laser exposure (∼10 mins.). Scale bars 20 μm.

The UV-resonance Raman spectra of single-phase solutions (**Figure 2a**, black curves) and phase-separated solutions (**Figure 2b**, black curves) of purified, unlabeled DDX4^[1-245]^ *in vitro* are similar, suggesting that DDX4^[1-245]^ does not undergo significant changes in secondary structure upon phase separation. Both single-phase and phase-separated solutions of DDX4^[1-245]^ show strong UV-resonance Raman bands at 1160, 1180, and 1205 cm^-1^ from phenylalanine (**Figures 2a-b**, green curves). The amide III and amide S (C_α_-H bending) modes that are sensitive to protein secondary structure appear between 1250-1350 cm^-1^ and at 1386 cm^-1^, respectively (**Figures 2a-b**, blue curves). The amide III mode at 1253 cm^-1^ is characteristic of conformations commonly seen in β-sheets, whereas the amide S mode at 1386 cm^-1^ can be present in both β-sheet and random coil configurations.^29^ An amide III vibrational mode at 1300 cm^-1^ is only expected for α-helix or β-sheet secondary structures, but not for random coil conformational behavior. The residuals (**Figures 2a-b**, purple curves) comparing alternative Fits 1 and 2 (**Figures 2a-b**, dashed red curves) to the experimental data (**Figures 2a-b**, black curves) show that the experimental spectra of single-phase and two-phase solutions have inferior fits in the absence of a vibrational mode at 1300 cm^-1^ (**Figures 2a-b**, bottom, dashed red curves). Therefore, the presence of an amide III vibrational band at 1300 cm^-1^ (**Figures 2a-b**, top, solid red curves) suggests DDX4^[1-245]^ deviates from total random coil conformational behavior, expanding on prior NMR studies.^25,26^

To probe the Raman response inside individual unlabeled DDX4^[1-245]^ condensates, we used confocal Raman microscopy (**Figure 2c**) with a visible laser (λ_excitation_ = 532 nm) and brightfield imaging (**Figure 2d**). The amide I vibrational mode, primarily C=O stretching of the protein backbone, of phase-separated DDX4^[1-245]^ condensates peaks at 1660 cm^-1^ (**Figure 2c**, black curve). The broad amide I vibrational response spanning 1600-1700 cm^-1^ could suggest contributions from α-helix and β-sheet secondary structures as well as random coil conformational behavior.^30^ However, the bending signal from H_2_O inside condensates also contributes throughout this same spectral region. Accurate assignment of the amide I Raman response thus requires deconvolution of spectral contributions from protein and the bending mode of water within condensates. Preparation of DDX4^[1-245]^ condensates in D_2_O shifts the spectral contribution of H_2_O out of the amide I region (**Figure 2c**, red curve), resolving the Raman contributions of arginine and aromatic tyrosine and phenylalanine sidechains ∼1550-1610 cm^-1^ and the amide I response that spans ∼1600-1690 cm^-1^ and peaks at 1660 cm^-1^. These observations are consistent with secondary structure conformational behavior that deviates from exclusively random coil conformations. Consistent with these spectroscopic results, all-atom simulations identified little persistent secondary structure, but found local deviations from self-avoiding random coil conformational behavior (**Figures S1 and S2**). Overall, our results using multiple approaches of UV-resonance Raman, confocal Raman microscopy, and computation each suggest that DDX4^[1-245]^ is disordered, yet possesses local conformational biases that deviate its backbone behaviors from those of a random coil.

Having investigated local conformational behavior using spectroscopic and computational methods, we next asked if the conformational behavior of monomeric DDX4^[1-245]^ under dilute conditions (*i*.*e*., the DDX4^[1-245]^ ensemble) possessed interactions that could cause the observed protein structural deviations away from a self-avoiding random coil. To examine this question, we turned to ensembles generated by STARLING, a deep-learning method for predicting conformational ensembles of intrinsically disordered regions.^31^ STARLING-derived ensembles predict a hydrodynamic radius of 41.3 Å, in good agreement with all-atom simulations (**Figure S3**), and reveal substantial intramolecular interactions (**Figure S4**). These results suggest that while DDX4^[1-245]^ is disordered, sequence-encoded interactions lead to deviations from self-avoiding random coil-like behavior.

To investigate the molecular interactions in the DDX4^[1-245]^ ensemble, we used STARLING to generate ensembles for previously characterized phase separation-deficient mutants: an all arginine-to-lysine variant (24RtoK, disrupting native cation-π interactions) and an all phenylalanine-to-alanine variant (14FtoA, disrupting cation-π and π-π interactions). Both ensembles reveal a substantial reduction of interactions (**Figure S4**), albeit with distinct impacts on the ensemble. They also showed a corresponding increase in hydrodynamic radius (*R*_*h*_) (**Figure S5**). We then generated an ensemble for a previously uncharacterized mutant that replaces all negatively charged residues with alanine (37EDtoA, disrupting electrostatic interactions), which again yielded an ensemble with a reduction of interactions and an increase in *R*_*h*_ compared to the wildtype sequence (**Figures S4-S5**). Taken together, we interpret these results to mean that intramolecular interactions in DDX4^[1-245]^ are driven by multiple distinct modes of interaction, including the π-π, cation-π, and electrostatic interactions.

### DDX4^[1-245]^ monomer collapse correlates with phase separation

To provide experimental support for our computational characterization, we next employed dynamic light scattering (DLS) to characterize monomeric DDX4^[1-245]^ under both non-phase-separating and phase-separating conditions. Specifically, we sought to understand how the global dimensions of DDX4^[1-245]^ change as a function of temperature, protein concentration, and ionic strength (**Figure 3**) when the system crosses the phase boundary. Nott *et al*., ^25^ Brady *et al*.,^26^ and Lin *et al*.,^27^ demonstrated that the DDX4 NTD undergoes homotypic phase separation to form condensates that are quantitatively described by physical models for phase separation. Moreover, this prior work established that homotypic phase separation of DDX4 NTD is suppressed at higher salt concentrations and temperatures.^24,25^ We therefore sought to understand how phase separation, as promoted or repressed by changes in parameters of temperature, protein concentration, and ionic strength, altered the conformational behavior of DDX4^[1-245]^.

**Figure 3.**
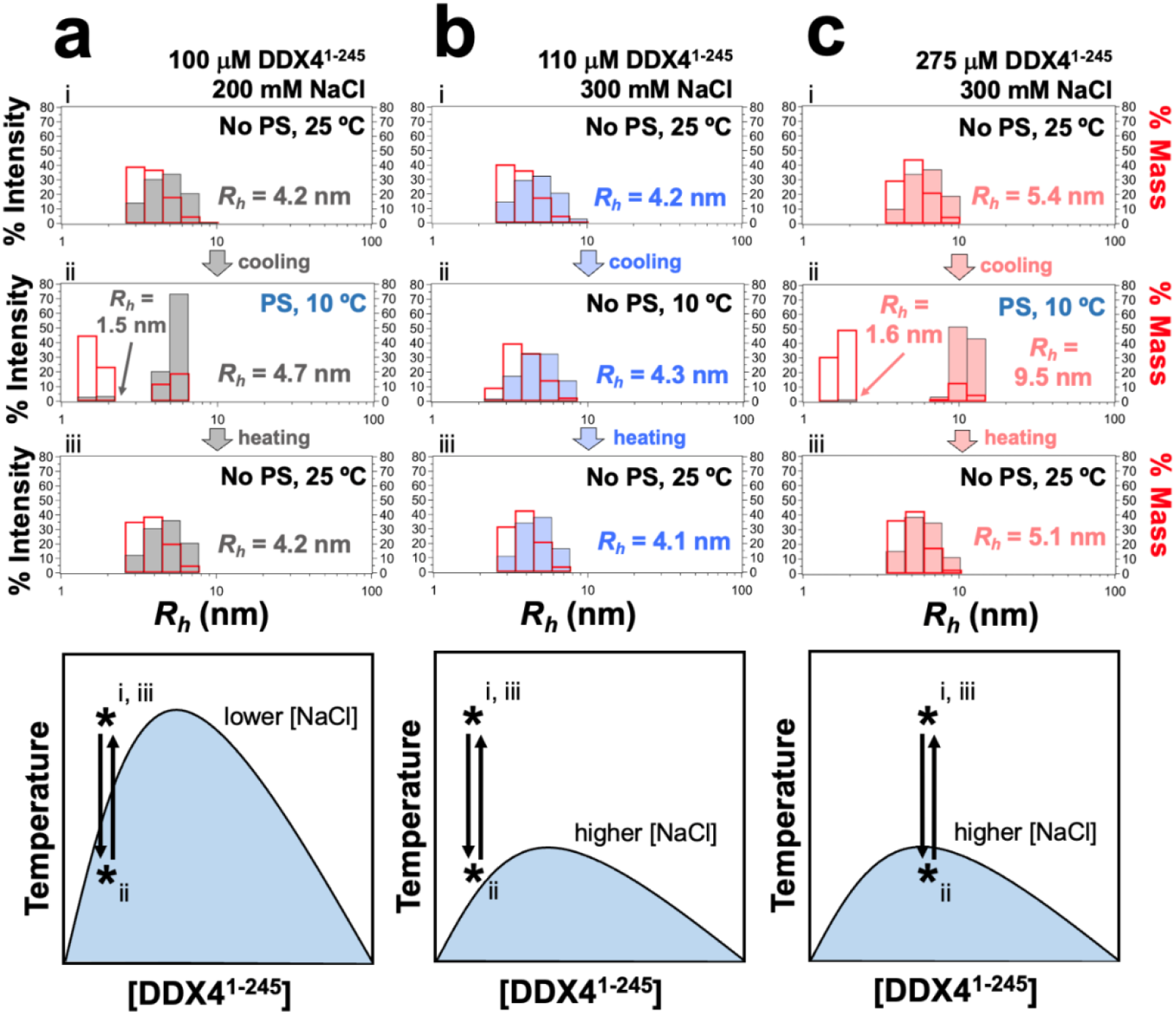
DDX4^[1-245]^ chain collapse correlates with phase separation. Representative dynamic light scattering measurements of monomeric DDX4^[1-245]^ as function of temperature, protein concentration, and NaCl concentration: % light scattering intensity of DDX4^[1-245]^ (gray, blue, and pink filled bars) are on the left axis, % mass of DDX4^[1-245]^ are on the right axis (red unfilled bars). Mean radius of hydration (*R*_*h*_) is reported for each observed population. Lower panels show schematic DDX4^[1-245]^ binodals (based on Brady *et al*.^26^) and corresponding experimental cycles of (i) starting point before cooling, (ii) stopping point after cooling, and (iii) end point after heating, with each point marked by an asterisk.

At a single-phase condition (100 µM DDX4^[1-245]^, 200 mM NaCl, 25° C), DLS reveals a single population of DDX4^[1-245]^ with apparent mean radius of hydration ⟨*R*_*h*_⟩ = 4.2 ± 1.1 nm (**Figures 3a and 3b**, top). This value is in good agreement with STARLING-derived ensembles with ⟨*R*_*h*_⟩ = 4.1 nm, supporting our computational and prior biophysical characterizations (**Figure S6**).^26^ However, the width of the distribution (±1.1 nm) suggests that dilute DDX4^[1-245]^ also forms oligomers.^32^ Indeed, under conditions in which phase separation is suppressed and protein concentration is increased (275 μM DDX4^[1-245]^, 300 mM NaCl, 10° C), the ⟨*R*_*h*_⟩ grows to 5.4 ± 1.3 nm. One possible interpretation of this result is that higher protein concentrations may favor formation of DDX4^[1-245]^ oligomers (**Figure 3c**, top) (see also *Discussion*).

Next, we sought to understand what happens to DDX4^[1-245]^ monomer under phase separating conditions. Phase separation of DDX4^[1-245]^ was induced by cooling the samples to 10 C (**Figures 3a-3c**, center). Surprisingly, under conditions at 10 °C where DDX4^[1-245]^ crossed the phase diagram binodal (**Figures 3a and 3c**),^25,26^ we observed a distinct protein population with dramatically reduced apparent ⟨*R*_*h*_⟩ = 1.6 ± 0.2 nm. We interpret this result to reflect compaction of DDX4^[1-245]^ monomer chains under phase separating conditions (**Figure S7**).

In principle, coil-to-globule chain collapse is expected at lower temperatures for polymers with an upper critical solution temperature (*i*.*e*., enthalpy-driven chain collapse)^33,34^ such as DDX4^[1-245]^ (**Figure S8**).^25^ We note that monomer collapse was not observed where DDX4^[1-245]^ is cooled under higher salt concentrations and does not cross the binodal (**Figure 3b**, center). Importantly, the collapsed monomer population also predominates under phase separating conditions, accounting for an estimated 70-80% of the sample by mass (% mass reported in **Figure 3** with red bars). Cycling the temperature to 25 °C and re-crossing the binodal shows that entry and exit of the monomer from this collapsed state were also reversible, a hallmark of phase separation. Thus, across conditions of temperature, protein concentration, ionic strength, and without the use of any molecular labels, we conclude from DLS that the appearance of a new, highly compact DDX4^[1-245]^ monomer population correlates strongly with DDX4^[1-245]^ phase separation.

### Dielectric constant is reduced inside DDX4^[1-245]^ condensates

Confocal Raman microscopy showed no significant structural changes of phase-separated DDX4^[1-245]^ in H_2_O or D_2_O (**Figure 2c**). However, intriguingly, under equivalent solution conditions, DDX4^[1-245]^ in D_2_O phase separates at a lower protein concentration than in H_2_O (**Figure 4a**), and phase separation in D_2_O also yields more condensates (**Figure 2d**). As DDX4^[1-245]^ exhibits no secondary structure changes in D_2_O versus H_2_O (**Figure 2c**), our observations suggest that water plays a key role in tuning the phase boundaries of DDX4^[1-245]^ (see also *Discussion*).^26,35^

**Figure 4.**
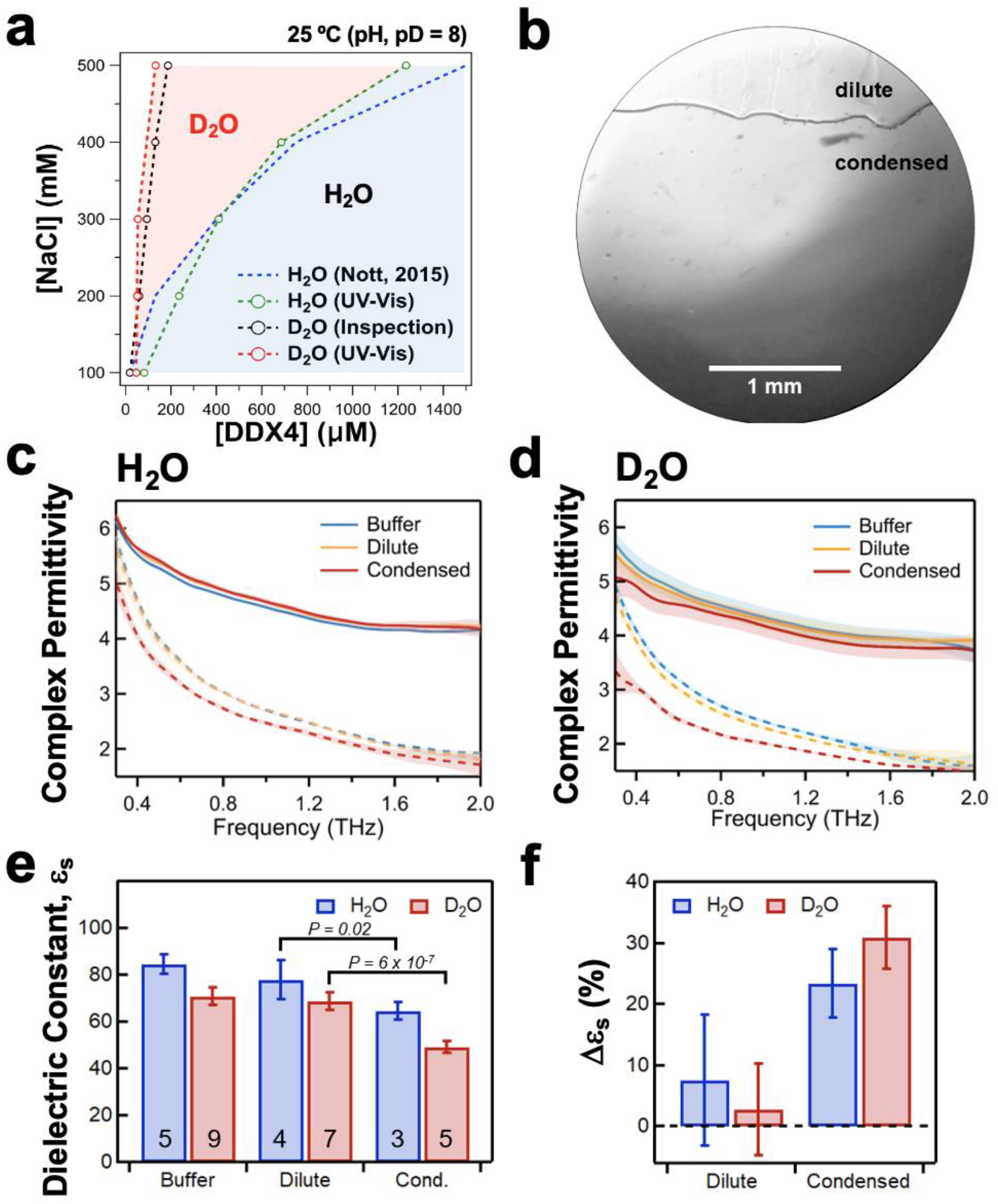
Terahertz-based quantification of the complex permittivity of monomeric and condensed DDX4^[1-245]^. (a) Phase diagram binodal of DDX4^[1-245]^ measured in (blue) H_2_O or (red) D_2_O, constructed based on Nott *et al*.^25^ (blue dashed curve), UV-visible absorbance of the dilute phase (green and red dashed curves, “UV-Vis”), or brightfield microscopy (black dashed curve, “Inspection”). (b) Spontaneous separation of DDX4^[1-245]^ in D_2_O into (top) dilute and (bottom) condensed phases within the terahertz (THz) cuvette. (c,d) Complex permittivity of buffer (blue), dilute (yellow), and condensed phases (red) of phase-separated DDX4^[1-245]^ prepared in H_2_O or D_2_O measured by terahertz (THz) time-domain spectroscopy. The shaded area represents the standard deviation across multiple measurements. (e) From (c,d), the static dielectric constant (ε_s_) of buffer, dilute, and condensed phases in H_2_O (blue) or D_2_O (red). The number of measurements collected for each condition is also shown. (f) The percent difference of ***ε***_***S***_ between dilute and condensed phases in H_2_O or D_2_O compared to buffer.

Using NMR, Brady *et al*. previously estimated that the static dielectric constant (ε_s_) of the *in vitro* DDX4^[1-236]^ condensed phase could range from 20 to 47 (as compared to 80 for pure H_2_O).^26^ However, the complex permittivity (also known as the dielectric response) inside condensates has never been directly quantified. Therefore, we aimed to directly measure the complex permittivity inside DDX4^[1-245]^ condensed phase using terahertz (THz) spectroscopy.

THz time-domain spectroscopy (THz-TDS) measures the phase-resolved transmission of coherent far-infrared radiation (∼1 THz = 33 cm^-1^), enabling the direct extraction of the frequency-dependent complex refractive index, and therefore permittivity, without requiring a Kramers-Kronig approximation (**Figure 1d**). In aqueous solutions, the dielectric loss in the THz region is dominated by Debye relaxation of water molecules aligned by the transient electric field. The phase-resolved transmission of THz radiation by water is used to extract the frequency-dependent complex refractive index (*n*) of the solution, which is equal to the square root of the complex permittivity (dielectric response, ***ε***) (**Figure 1d**).^36^ The resulting dielectric spectrum can then be fit to a constrained Debye model to semi-quantitatively extract the static permittivity (*i*.*e*., dielectric constant, ***ε***_***S***_). After inducing phase separation, we concentrated DDX4^[1-245]^ via centrifugation in H_2_O and D_2_O to aquire dilute and condensed layers, then carefully transferred each layer to a quartz microcuvette by pipetting (**Figure 4b**) and measured the complex-valued permittivity spectra (0.25-2.0 THz) of the dilute phase, the condensed phase, and the buffer for both H_2_O and D_2_O (**Figures 4c and 4d**).

In both H_2_O and D_2_O, the imaginary component of the complex permittivity (**Figures 4c and 4d**, dashed lines) was suppressed in the DDX4^[1-245]^ dense phase. Fitting the data to a constrained double Debye model (see Methods and **Tables S1-S2**) gave static dielectric constants of pure buffer in the expected range (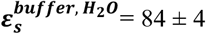 and 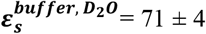), whereas the condensed phase static dielectric constant (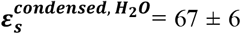 and 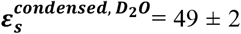) was significantly reduced relative to the dilute phase (**Figure 4e**). For buffer solutions, our static dielectric constant is overestimated by ∼7% for H_2_O based on tabulated values (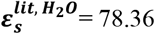 at 25 °C), but is underestimated by ∼9% for D_2_O (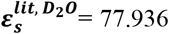 at 25 °C).^37^ Because THz-TDS measures the “high-frequency” dielectric response, measurement of the static dielectric constant relies on changes in the amplitudes and lifetimes of the Debye relaxation terms used in the fitting model. Relative to buffer, the percent change of static dielectric in the condensed phase corresponded to a change of ∼20% in H_2_O and ∼30% in D_2_O (**Figure 4f**), indicating that our measurement is still sensitive to the relative change in static dielectric constant upon condensation at a semi-quantitative level. Overall, our analysis suggests that although 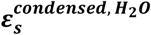 is substantially lower than in the dilute phase, the reduction of 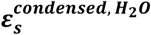 is less significant than previously predicted.^25,26^ In sum, relative to the surrounding bulk aqueous environment, condensates offer unique chemical environments. Moreover, the solvent itself can have a major impact on the intracondensate chemical environment, as seen here for H_2_O versus D_2_O.

### Optical observation and quantification of reduced water content inside DDX4^[1-245]^ condensates

Prior NMR experiments suggested ∼75% volume fraction of water inside DDX4^[1-236]^ condensates.^26^ We aimed to directly image and quantify the reduced water content inside DDX4^[1-245]^ condensates using stimulated Raman scattering (SRS) imaging (**Figure 5**). Whereas conventional spontaneous Raman scattering emits relatively few inelastically-scattered photons, SRS can achieve an enhancement factor as high as ∼10^8^ over spontaneous Raman.^38^ Our SRS imaging achieves point scanning with a lateral resolution of ∼330 nm and an axial resolution of ∼1.4 μm.

**Figure 5.**
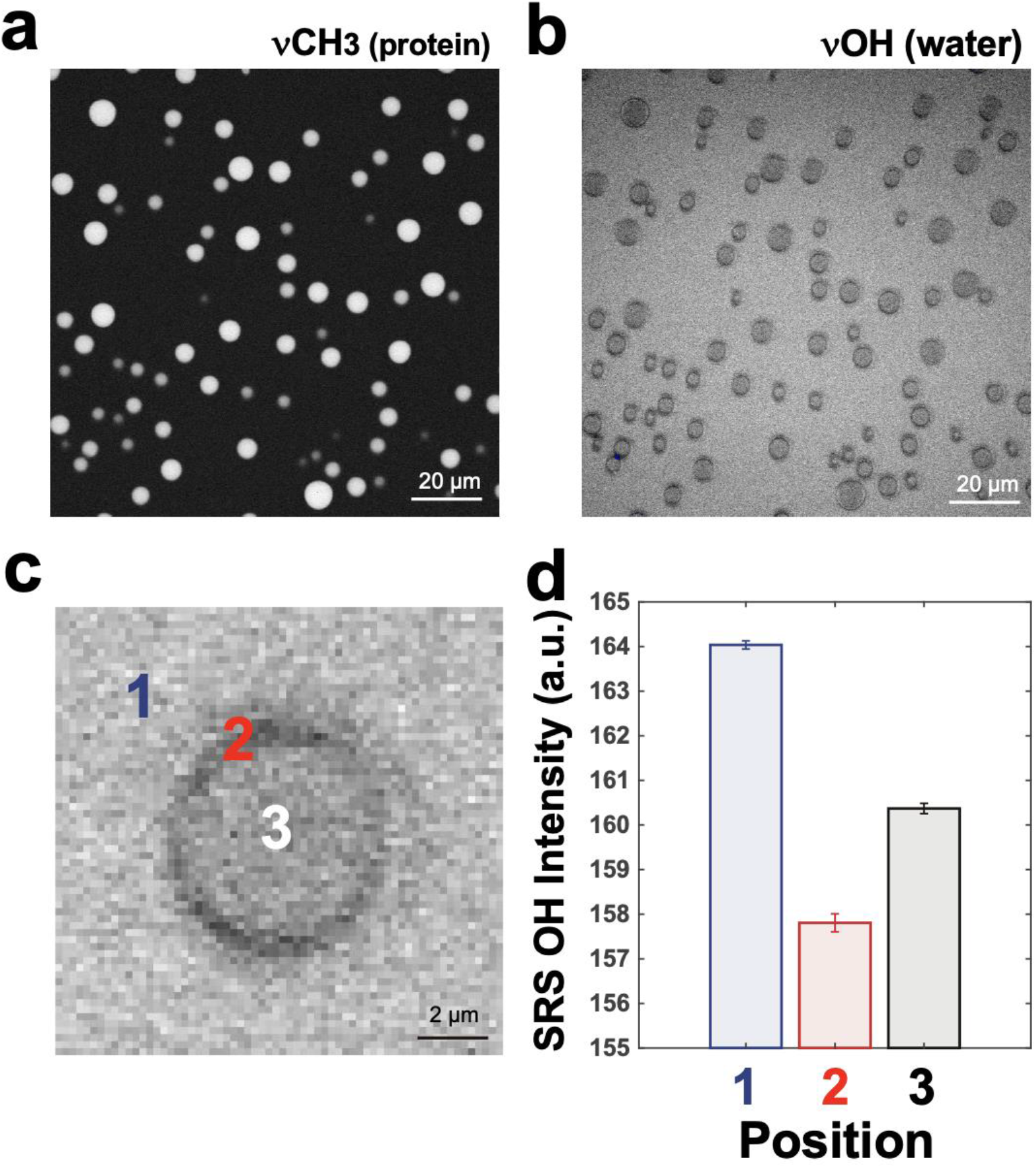
Stimulated Raman Scattering (SRS) quantification of water release from DDX4^[1- 245]^ condensates. (a) SRS imaging (λ_Stokes_ = 1064.2 nm, λ_pump_ = 810.5 nm) detects CH_3_ vibrational response of DDX4^[1-245]^ protein concentrated in condensates. (b) SRS imaging (λ_Stokes_ = 1064.2 nm, λ_pump_ = 790 nm) reveals reduced O-H vibrational response of water inside and at the interface of DDX4^[1-245]^ condensates. (c) Detail of an individual condensate O-H vibrational response imaged with SRS. (d) Quantification (mean ± SEM) of SRS water O-H intensity at positions (1) outside, (2) at the interface, and (3) inside condensates for *N* = 67 condensates.

We imaged DDX4^[1-245]^ condensates in H_2_O using SRS of the CH_3_ stretching (**Figure 5a**) and the O-H stretching (**Figure 5b**) vibrational modes. The O-H stretching showed a strong SRS response both inside and outside the condensates because the overwhelming contribution was due to water (**Figure 5b**). However, compared to the dilute phase, DDX4^[1-245]^ condensates appear darker in SRS images due to their lower water content (**Figure 5b**). Within condensates, SRS imaging revealed even lower water content at the interface of condensates compared to the condensate core (**Figure 5c**). SRS intensity differences of water O-H stretching showed that water content within the condensate core was ∼2.5% lower compared to bulk, while water content at condensate perimeters was ∼4% lower (**Figure 5d**). Thus, water content inside individual DDX4^[1-245]^ condensates is reduced compared to bulk. However, the measured water content is much higher than previously estimated at ∼75% for a collected DDX4^[1-236]^ condensed phase.^26^

## DISCUSSION

Biomolecular condensates are non-stoichiometric assemblies that, in many cases, are well described by the physics of phase separation. Condensates can potentially enable the design of novel and tunable chemical environments. For decades, industrial chemists have sought to refine ideal solvents for complex chemical pathways. In the same vein, one possible role for biomolecular condensates is to form novel intracellular solvent environments, enabling or catalyzing specific biochemical reactions. Investigating this concept necessitates the ability to probe intracondensate chemistry, yet this has been historically challenging to accomplish in a non-perturbative manner.

Optical spectroscopies with imaging capabilities provide spatially resolved, label-free tools for investigating the chemistry of biomolecular condensates. Several publications have employed Raman and attenuated total reflection Fourier transform infrared (ATR-FTIR) spectroscopies for the determination of protein structure^39-41^ and protein concentration^42,43^ in biological condensates. Surface-enhanced Raman spectroscopy (SERS) using silver nanoparticles was employed to investigate the role of arginine, tyrosine, and tryptophan residues in the formation of cation-π and π-π interactions driving the phase separation of Fused in Sarcoma (FUS) condensates.^44^ ATR-THz spectroscopy of FUS and other protein condensates suggests that release of entropically unfavorable hydration around hydrophobic protein surfaces could promote condensate formation, while retention of enthalpically favorable hydration at hydrophilic protein surfaces can help maintain condensates in a liquid-like state.^19-23,45,46^ Two-dimensional infrared (2D-IR) spectroscopy of heterotypic poly-arginine/adenosine monophosphate condensates showed that water dynamics are significantly slowed within condensates due to strong electrostatic interactions.^47^ Consistent with these findings, hyperspectral imaging of solvatochromic dyes suggested lower dipolar relaxation of water inside glycinin protein condensates.^41^ Intracellular quantitative refractive index (RI) imaging was also explored for detection of nucleoli, heterotypic nuclear speckles and cytoplasmic stress granules,^48^ as well as the internal structure of synthetic single-component ferritin-based condensates,^49^ although the lack of contrast of intracellular cytoplasmic stress granules and nuclear speckles in RI imaging suggests these condensate densities are comparable to the crowded, aqueous cellular environment. More recently, a label-free method was developed that integrated quantitative phase imaging with refractive index measurements to deconvolve compositions of multi-component condensates, enabling detailed experimental characterization of multi-component phase diagrams.^50^

In the present work, we applied UV-resonance Raman, confocal Raman, stimulated Raman scattering (SRS), and terahertz (THz) spectroscopies in combination with dynamic light scattering (DLS), as well as computational ensemble prediction methods to probe the conformational behavior of individual proteins and the chemical and physical properties of condensate interiors and interfaces. Our application of this suite of complementary methods has enabled quantifying the critical roles of protein hydration and protein collapse, leading to a proposed molecular model underlying DDX4^[1-245]^ phase separation (**Figure 6**).

**Figure 6.**
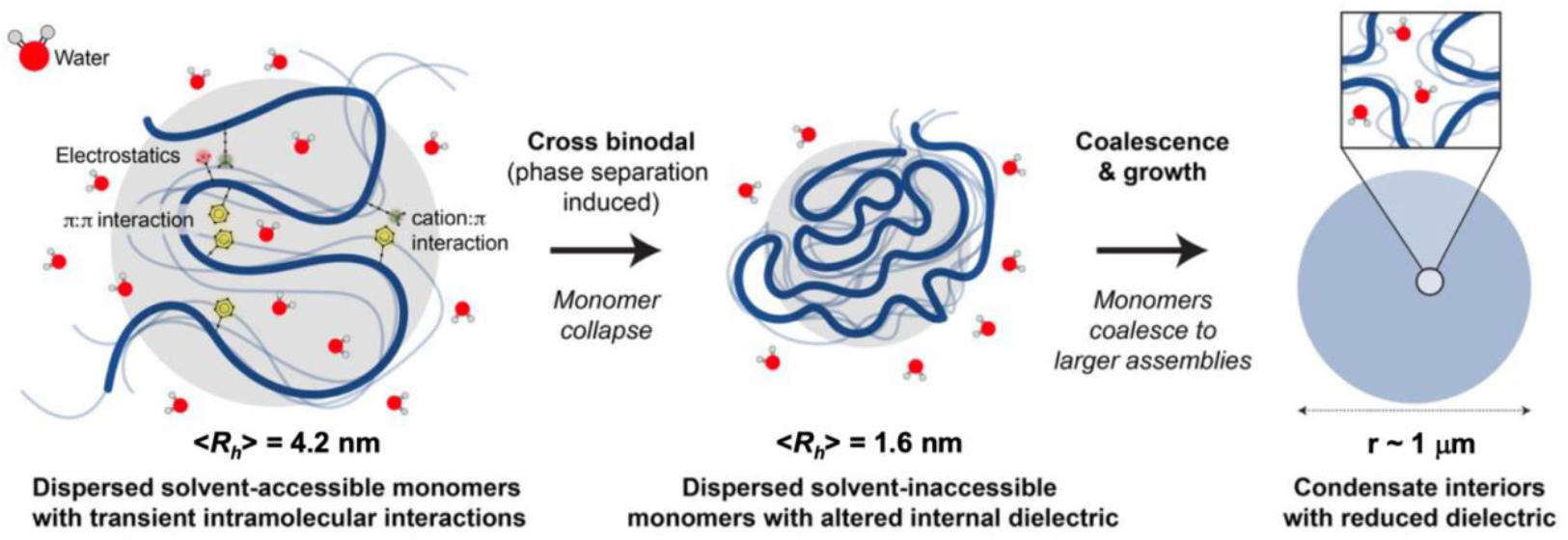
Model of DDX4^[1-245]^ liquid-liquid phase separation. Our results from terahertz spectroscopy, stimulated Raman scattering imaging, and dynamic light scattering suggest that phase separation (PS) triggers DDX4^[1-245]^ protein chain collapse, water release, and reduction of the static dielectric constant (***ε***_***S***_). A reduced dielectric and water release increase the strength of protein-protein arginine-phenylalanine cation-π interactions inside condensates, driving condensate growth and phase separation, possibly via positive feedback.

For one-phase solution conditions, DDX4^[1-245]^ exists as monomers that can form small oligomers at high protein concentration. Using DLS, we detect a single broad population with apparent mean radius of hydration ⟨*R*_*h*_⟩ = 4.2 ± 1.1 nm (**Figures 3a and 3b**), which increases with increasing protein concentration (**Figure 3c**). This hydrodynamic radius is in good agreement with ensembles generated using STARLING (⟨*R*_*h*_⟩ = 4.1 nm) (**Figure S6**). STARLING-derived ensembles for DDX4^[1-245]^ suggest attractive interactions driven by cation-π, π-π, and electrostatic interactions. Such interactions cause deviations in the single-chain ensemble away from the behavior expected for a real chain with excluded volume but no meaningful attractive or repulsive interactions (*i*.*e*., a self-avoiding random coil). Moreover, DDX4 variants deficient in phase separation show a substantial diminution in intramolecular interactions, consistent with prior work.^9,51,52^

The observation of chain deviations from self-avoiding random coil conformational behavior is consistent with our analysis of UV-resonance Raman amide III response (**Figures 2a and 2b**). Prior NMR work on the DDX4 NTD excluded the possibility of the stable long-lived acquisition of secondary structure.^26^ Nevertheless, our UV-resonance Raman amide III response is incompatible with a conformational ensemble that is exclusively random coil, suggesting that non-random intramolecular interactions give rise to conformations that deviate from random coil configurations. In addition, UV-resonance Raman probing the amide III response (**Figures 2a and 2b**) and confocal Raman microscopy probing the amide I response (**Figure 2c**) both suggest that DDX4^[1-245]^ molecules retain their dilute-phase conformational biases in the condensed phase.

An important aspect of our model proposes that DDX4^[1-245]^ monomers can undergo collapse (⟨*R*_*h*_⟩ = 1.6 ± 0.2 nm) in conditions in which phase separation can occur *(e*.*g*., reduced temperature, reduced salt concentration) (**Figure 3 and Figure S9**). For polymers with upper critical solution temperature, such as DDX4^[1-245]^, phase separation is expected to be enthalpy-driven and chain collapse is expected.^25,33,34^

In addition to monomeric DDX4^[1-245]^, we also observe the appearance of larger species by DLS (⟨*R*_*h*_⟩ = 5.4 nm ± 1.3 nm) even under conditions in which phase separation is suppressed (**Figure 3c**). One simple explanation for these larger species is that they are small oligomers (estimated to be dimers or trimers) of DDX4^[1-245]^ formed from monomeric DDX4^[1-245]^ that retain dimensions of around 4.2 ± 1.1 nm. An alternative interpretation is that these larger species are clusters of tens of highly compact DDX4^[1-245]^ monomers, where each monomer in the cluster has an *R*_*h*_ closer to the compact population observed under phase separating conditions. Such an interpretation could be consistent with prior work in which clusters of disordered proteins were observed under subsaturated conditions.^32^ However, even in that prior work, a small increase in the dimensions of monomers was interpreted as small oligomers, whereas clusters were found to be an order of magnitude larger than monomers. As such, these findings currently favor the simpler explanation of small oligomers. Further studies using label-free high-resolution mass photometry^53^ could potentially help to resolve these possibilities.

The DDX4^[1-245]^ collapsed monomer can be considered a single chain condensate, in other words, the smallest possible condensate unit. The parallels between single-chain condensation and multi-chain phase separation have been drawn since the 1970s.^54-56^ For synthetic polymers, single-chain collapse across conditions that mirror the phase boundary was also reported.^57^ Recently, time-resolved SAXS revealed single-chain compaction of the low-complexity prion-like domain from the RNA-binding protein hnRNPA1 upon transition into phase-separation-compatible conditions and prior to coalescence and growth.^58^ This work focused on the kinetics of phase separation, and suggested this compact species was relatively transient. In contrast, here we observe a population of compact monomers that persists for at least timescales accessible to DLS without the need for a stopped-flow or microfluidics apparatus. As such, we propose that the DDX4^[1-245]^ collapsed state may represent an experimentally accessible target for studying early-stage condensate nucleation and growth. Further characterization of the DDX4^[1-245]^ collapsed state, especially via single-molecule techniques, may offer a powerful model system to dissect the specific molecular interactions that underlie the kinetics, thermodynamics, and chemical driving forces of condensate nucleation and growth.

As reported previously, DDX4^[1-245]^ is overrepresented in phenylalanine and arginine residues compared to other IDPs, and cation-π interactions appear to be one important mode of intermolecular interaction that underlies its phase behavior.^1,24-26^ STARLING ensembles support the importance of these interactions, and show that losing either native arginine residues (24RtoK mutant) or native phenylalanine residues (14FtoA mutant) leads to an increase in global dimensions and loss of intramolecular interactions (**Figures S4-S5**). Moreover, our work predicts that the loss of acidic residues will also suppress intramolecular interactions, reducing homotypic phase separation (**Figures S4-S5**). The importance of arginine-aromatic interactions in driving condensate formation for intrinsically disordered regions is well-established, with arginine-to-lysine subsistutions shown in many contexts to abrogate attractive interactions essential for phase separation.^8,25,26,39,59-61^ Prior work suggests that arginine-phenylalanine interactions are stronger than lysine-phenylalanine interactions.^62^ Computational work has indicated that a major determinant of arginine’s importance (compared to lysine) stems from the difference in free energy of hydration between the residues.^63^ The calculated free energy of hydration difference (∼12 kcal/mol) between positively-charged arginine and lysine residues shows that arginine is more easily dehydrated, consistent with a model in which protein dehydration and water release from arginine-rich condensates can help to promote phase separation. Therefore, besides protein chain collapse, any proposed model must also account for how water behavior may drive DDX4^[1-245]^ phase separation.

Our development of THz-TDS and *in situ* stimulated Raman spectroscopy of condensed phase DDX4^[1-245]^ (**Figures 4-5**) has enabled us to directly probe water release and reduced static dielectric, which can drive phase separation and tune protein-protein interactions inside condensates. Inside the DDX4^[1-245]^ condensed phase, the dielectric is reduced to 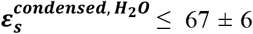 and 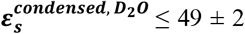 (**Figure 4e**). The water content ranges from ∼97.5% at the condensate core to ∼96% at the condensate interface (**Figure 5d**). The significant reduction of the dielectric response in the condensed phase that we observe (**Figure 4**) cannot fully be explained by the measured water release (**Figure 5**). Therefore, the reduction of the dielectric response can only be explained if the dielectric response of water in close proximity to DDX4^[1-245]^ differs markedly from the dielectric response of pure water. The high dielectric value of free water is related to the free rotation of the static dipoles of water molecules in an external field. Our observation of reduced dielectric response of water inside condensates strongly suggests that these water molecules are not freely rotating and form either a solvation shell or a caged geometry, consistent with recent reports from 2D-IR spectroscopy of water dynamics inside condensates.^47^ In both cases, the polarizability per molecule will be lower, resulting in the reduced overall dielectric response.

Cation-π protein-protein interactions are believed to help drive DDX4^[1-245]^ phase separation. In the aqueous phase dominated by the high dielectric constant of H_2_O (∼80), the cation-π interaction energy is similar to a single hydrogen bond (∼5 kcal/mol), but becomes favorable (∼19 kcal/mol in the gas phase) as the dielectric of the medium is reduced.^64,65^ Computational predictions suggest that our experimentally observed changes in dielectric inside DDX4^[1-245]^ condensates (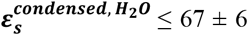 and 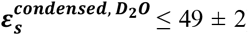) would increase the interaction strength of individual cation-π interactions by <0.1 kcal/mol,^65^ which emphasizes the importance of multivalent interactions in driving biological phase separation. As condensates form and grow, increasingly stronger protein-protein interactions may promote further water release and reduction of the dielectric, likely establishing positive feedback that can drive phase separation and condensate growth. Significantly, our experimental data suggest a model that unifies the concepts of protein chain collapse, reduced dielectric, and protein dehydration as mechanisms that strengthen multivalent protein-protein interactions to drive phase separation.

We also observe that D_2_O significantly shifts the DDX4^[1-245]^ binodal compared to H_2_O (**Figure 4a**). The impact of D_2_O on phase separation has been reported previously in the context of a dehydrogenase,^66^ lysozyme,^67^ aprotinin,^68^ γB-crystallin,^69^ bovine serum albumin,^70^ and androgen receptor proteins,^71^ where qualitatively similar observations were reported, but the origin of this isotope effect remains unclear. Neutron scattering studies of protein solubility and phase separation^66-70^ observed that attractive protein-protein interactions are strengthened in D_2_O, although the physical mechanisms (*e*.*g*., stronger hydrophobic or electrostatic interactions) underlying this phenomenon remain debated. Our observation that 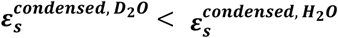 (**Figure 4**) likewise implies stronger attractive protein-protein interactions. Besides protein-protein interactions, D_2_O hydrogen bonds are also stronger by 0.1-0.2 kcal/mol compared to H_2_O.^72^ Both stronger protein-water interactions involving hydrogen-bonding interactions with D_2_O and stronger protein-protein interactions will enthalpically (**Δ*H***^***Protein***−***water***^ and **Δ*H***^***Protein***−***Protein***^) favor phase separation, counterbalancing the free energy costs of reduced numbers of hydration water molecules and protein configurational entropy (**Δ*S***^***Protein***−***water***^ and **Δ*S***^***Protein***−***Protein***^). These conditions are consistent with DDX4^[1-245]^ phase separation being enthalpy-driven, and suggest that D_2_O may also promote DDX4^[1-245]^ chain collapse to drive phase separation.

## CONCLUSIONS

Our work demonstrates that label-free vibrational spectroscopies and optical scattering methods provide new insights into the underlying physical chemistry associated with condensate formation. Through integration of light-scattering and computational methods, we identify a previously unreported collapsed monomeric species that strongly correlates with phase separation. Our measurements quantify critical physical, chemical, and structural properties of phase separated condensates that until now have proven experimentally refractory and computationally inaccessible. Our data leads us towards a model of phase separation that unifies the molecular concepts of protein dehydration, water release, reduced water polarizability, protein chain collapse, and protein-protein interactions as critical driving forces of homotypic protein phase separation. Further development of the optical methods demonstrated here should enable fundamental investigations into homotypic and heterotypic condensate interiors and interfaces, enabling greater understanding of the relationships between protein charge, enzymatic activity, and the chemical and physical properties of biological microenvironments. Our insights will inform the design of custom chemical microenviroments that can be tailored for greater control of chemical reactions.

## METHODS

### Expression and Purification of DDX4^[1-245]^

DDX4^[1-245]^ was recombinantly expressed in *E. coli* with an N-terminal GST domain and TEV-protease cleavage site. From glycerol stock, bacteria inoculates were grown in LB containing 0.1 mg/mL ampicillin at 37 °C overnight. The inoculates were transferred to 1 L of LB with ampicillin and grown at 37 °C until OD_600_ = 0.6-0.8 AU. The cultures were then induced with 0.25 mM IPTG. The cultures were immediately brought to room temperature and grown overnight. Cells were collected by centrifugation and resuspended in lysis buffer (50 mM Tris, 500 mM NaCl, 5 mM DTT, pH = 8) containing protease inhibitor cocktail (cOmplete Mini, EDTA-free, Sigma-Aldrich, Cat. No. Roche-11836170001). Cells were sonicated on ice three times for 7 minutes each time in steps of 3 seconds (followed by 5 seconds off). The sonicate was clarified by centrifugation.

GST beads (Glutathione Sepharose 4B, GE Life Sciences, Cat. No. 17-0756-01) were washed three times with 5 mL binding buffer (500 mM NaCl, 10 mM Na_2_HPO_4_, 1.8 mM KH_2_PO_4_, pH = 7.3) in 15 mL Falcon tubes (1 mL of beads per 1 L of cell culture, using 1 mL beads per Falcon tube). Beads were subsequently washed once in 5 mL lysis buffer. The buffer was removed, and approximately 12 mL of clarified sonicate was applied per 1 mL of GST beads. The Falcon tubes were nutated overnight at 4 °C to ensure complete binding of protein to beads.

Beads were decanted by centrifugation at 4 °C and washed twice with 5 mL lysis buffer per 1 mL beads. TEV protease solution (20 uL ProTEV Plus, Promega, Cat. No. V6101, in 980 μL lysis buffer) was applied to 1 mL of GST-TEV-DDX4^[1-245]^-bound beads. Beads were incubated at room temperature for 5 hours while nutating. The supernatant containing TEV protease and DDX4^[1-245]^ was decanted by centrifugation at room temperature. Beads were washed with 1 mL lysis buffer, centrifuged, and the supernatant collected to recover remaining protein.

FPLC purification (Superdex 75 prep grade, GE Life Sciences) was performed in storage buffer (500 mM NaCl, 20 mM Tris, 5 mM TCEP, pH 8 in H_2_O) and used to separate TEV protease (48 kDa) from DDX4^[1-245]^ (26.7 kDa). This yielded approximately 9-10 mg of DDX4^[1-245]^ per liter of bacterial culture. Purified DDX4^[1-245]^ was verified by SDS-PAGE, which showed a single band at ∼26 kDa. DDX4^[1-245]^ was concentrated to 500-600 μM (Amicon Ultra, 3 kDa cutoff, Millipore, Cat. No. UFC900324). Aliquots were flash frozen in liquid nitrogen and stored at -80 °C until use. Aliquots were thawed at room temperature for at least 30 minutes prior to use in all experiments.

### UV-Resonance Raman Spectroscopy

FPLC-purified DDX4^[1-245]^ in storage buffer was quickly diluted (“quenched”) at room temperature to a final concentration of 50 μM DDX4^[1-245]^ and 125 mM NaCl (with 20 mM Tris, 5 mM TCEP, pH = 8 in H_2_O). This condition yielded phase-separated condensates while avoiding the growth of larger condensates over the duration of the experiments. The sample was probed at room temperature (25 °C) using a home-built UV-resonance Raman spectrophotometer. Briefly, the 215 nm light output from the fourth harmonic of a 1 kHz Ti:Sapphire laser was focused on a vertically-mounted, fused silica microcapillary with inner diameter 100 µm. The protein sample was flowed at a rate to ensure fresh sample for each pulse. The Raman-scattered photons were collected, dispersed through a prism prefilter and spectrograph, and detected by a CCD. The power at the sample was less than 0.8 mW. Five 1-minute spectra of protein sample and buffer were collected to generate the final UV resonance Raman difference spectra of protein.

### Confocal Raman Spectroscopy

FPLC-purified DDX4^[1-245]^ was quenched at room temperature to a final concentration of 115 μM in 100 mM NaCl, 20 mM Tris, 5 mM TCEP, pH = 8 (pD = pH + 0.4 = 8) in H_2_O (or D_2_O), yielding phase-separated condensates. The phase-separated mixture was applied to a glass slide (Fisher Scientific, Cat. No. 12-544-4) and sealed with a glass coverslip (Fisher Scientific, Cat. No. 12-545-A) using a spacer (Sigma-Aldrich, Grace Bio-Labs SecureSeal Imaging Spacer, GBL654008). The sample was probed at room temperature using a home-built confocal Raman microscope. A 532-nm laser (Samba 532 nm, 400 mW, Cobolt Inc.) is used as the light source. The laser beam was first collimated and expanded by telescope lenses (f_1_ = 100 mm, f_2_ = 300 mm, Thorlabs) to achieve the designed illumination spot size. The expanded beam was then directed to an inverted microscope (IX71, Olympus) with a dichroic beam splitter installed (LPD1, LPD02-532RU-25, Semrock). A UplanSApo 20×, N.A. = 0.75 (Olympus) objective was used. The emitted Raman signal first passed a pinhole (100 µm, Thorlabs) for background suppression and was relayed by two lenses (Thorlabs) before being projected to the spectrometer (Kymera 328i, Andor) that is equipped with a grating of 600 lines/mm, blazed at 500 nm. A long-pass filter (LP03-532RU-25, Semrock) was installed between the relay lenses to block laser light from Rayleigh scattering. Raman signal was then collected by an EMCCD with an active pixel-area of 200×1600 (Newton970, Andor). All the data collection was performed with a custom LABVIEW program (National Instruments). Using a laser power of 0.1 mW, spectra were collected by moving the laser beam inside or outside of condensates. No laser-induced changes of condensates were observed inside of condensates. When measuring outside of the condensates, small condensates could become optically trapped in the path of the laser, although this occurred on timescales longer than our experiments.

### Dynamic Light Scattering

FPLC-purified DDX4^[1-245]^ in storage buffer was filtered (GE Healthcare Life Sciences, Whatman Anotop 10 mm syringe filter, 0.1 μM pore size, Cat. No. 6809-1012) at room temperature and prepared at the concentrations of DDX4^[1-245]^ and NaCl described in the main text (all conditions at 20 mM Tris, 5 mM TCEP, pH = 8). Experiments were carried out using a Wyatt DynaPro PlateReader II instrument. Sample solutions (50-100 μL) were transferred to wells in a 384-well glass-bottom plate (Greiner Bio-One SensoPlate, VWR Cat. No. 82051-546). For each experiment, 60-100 acquisitions were collected, 5 seconds per acquisition, and the quality of the autocorrelation function was verified. The instrument was cycled from 25 C to 10 °C to 25 °C, and measurements were performed on the same sample at these temperatures. The experimental data were analyzed in the provided Dynamics software (Wyatt Technology) using the “linear polymers” protein model, 2% sodium chloride buffer, and a cutoff of %Mass ≥ 0.5%.

### Terahertz Spectroscopy

Terahertz time-domain spectroscopy (THz-TDS) measurements were performed using a home-built spectrometer as previously described in detail elsewhere.^36^ Briefly, the output of a Ti:Sapphire oscillator (Spectra-Physics Mai Tai SP, 800 nm center wavelength, 84 MHz repetition rate, 3.5 nJ pulse energy, <35 fs pulse duration) was split into two beams for generation and detection of THz radiation using two separate photoconductive antennae (Batop). The time delay between generation and detection beams were controlled using a mechanical delay stage, allowing the transient electric field to be measured.

Samples were centrifuged through the onset of PS and prepared for measurement by pipetting ∼10 μL of condensed phase, dilute phase, or storage buffer solution into the bottom portion of a 100 μm pathlength, two-piece quartz cuvette (Starna Cells). The two-piece cuvettes were sealed with clear nail polish to prevent evaporation of the sample. The sample cuvette was mounted on a holder attached to a mechanical positioning stage, which allowed for the sample measurement, reference measurement through an empty portion of the cuvette, and an air reference measurement to be collected sequentially. The complex refractive index (*n*) of measurements was processed numerically as previously described^36^ and converted to the complex permittivity (ε), or dielectric spectrum, using the relation ***ε***_***S***_ = ***n***^**2**^, which holds for non-magnetic materials.

The resulting permittivity spectra were fit to a constrained double Debye model as shown below in Eq. 1 where ***ε***(***ω***) is the frequency-dependent permittivity, ***ω*** is the frequency, ***ε***_∞_ is the dielectric constant in the high-frequency limit, **Δ*ε***_***i***_ is the relaxation amplitude of the *i*^th^ Debye resonance (*i* = 1, 2), and ***τ***_***i***_ is the relaxation time of the *i*^th^ Debye resonance. For the final term corresponding to the intermolecular stretching vibrational mode, ***A***_***S***_ is the amplitude, ***ω***_***S***_ is the angular frequency, and ***γ***_***S***_ is the damping constant.

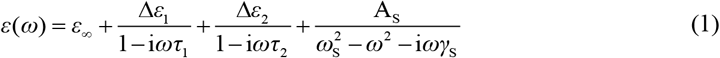

Because the first Debye resonance (**Δ*ε***_**1**_) is below the bandwidth of the spectrometer used herein, ***τ***_**1**_ and ***τ***_**2**_ cannot be accurately fit simultaneously. Therefore, the low-frequency Debye resonance lifetime (***τ***_**1**_) was fixed to 8.76 ps for both H_2_O and D_2_O based on previous THz-TDS measurements on H_2_O in the literature.^73^ Likewise because the intermolecular stretching vibrational mode is higher in frequency than the bandwidth of the spectrometer used herein and has a significant contribution to the total dielectric constant, we fixed ***ω***_***S***_/**2*π*** (5.30 THz and 5.36 THz for H_2_O and D_2_O, respectively) and ***γ***_***S***_/**2*π*** (5.40 THz and 5.06 THz for H_2_O and D_2_O, respectively) based on literature values.^74^ The static dielectric constant (***ε***_***S***_) was then calculated using the obtained amplitudes from the fit according to Eq. 2 as shown below:

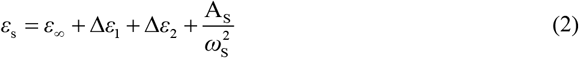

### Stimulated Raman Scattering (SRS) Imaging

FPLC-purified DDX4^[1-245]^ was quenched at room temperature in 100 mM NaCl, 20 mM Tris, 5 mM TCEP, pH = 8. The sample was probed at room temperature using a home-built SRS microscope. SRS uses a laser fixed at the Stokes wavelength (λ_Stokes_) and a laser tuned to the pump wavelength (λ_pump_) when the difference 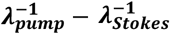 is close to the resonant vibrational frequency.

Two synchronized 6-ps lasers (called pump and Stokes beams) with 80-MHz repetition rate are provided by a picoEmerald system from APE (Applied Physics & Electronic, Inc.). The pump beam is tunable from 720-990 nm; the Stokes beam is fixed at 1064.2 nm. The intensity of the Stokes beam was modulated sinusoidally by a built-in electro-optic modulator (EOM) at 8 MHz with a modulation depth of more than 90%. Spatially and temporally-overlapped pump and Stokes beams were coupled into the laser scanning microscope. After passing through the specimens, forward-going pump and Stokes beams were collected with an IR-coated oil condenser (1.4 NA, Olympus). The Stokes beam is completely filtered with two high-optical-density bandpass filter (890/220 CARS, Chroma Technology) and the transmitted pump beam was detected by a large-area (10 mm × 10 mm) Si photodiode (FDS1010, Thorlabs). The output current of the photodiode was then sent to a fast lock-in amplifier (HF2LI, Zurich Instruments) for signal demodulation. The laser power was set as *P*_pump_=17 mW, *P*_Stokes_=67 mW; and images were generated with the pixel dwell time of 4 μs, and the time constants of the lock-in amplifier set to 2 μs.

### STARLING ensemble generation for DDX4^[1-245]^

STARLING ensembles were generated using STARLING version 1.0.0 (obtained April 2025) using the VAE/U-Net implementation with DDPM weights model-kernel-epoch.47-epoch_val_loss.0.03.ckpt and VAE weights model-kernel-epoch.99-epoch_val_loss.1.72.ckpt.^31^ STARLING is a deep-learning generative model trained on data from the Mpipi-GG forcefield (a derivative of the Mpipi forcefield) that enables the rapid generation of coarse-grained ensembles of disordered proteins in minutes on commodity hardware.^75^ Ensembles contain 6,000 conformers and were generated on a MacBook Pro (M3) using the command: `starling <sequence> -b 400 -c 6000` (note that the `-b` option here defines the batch size, which has no impact on the generated ensemble but scales the number of conformers generated in parallel). Ensemble generation takes ∼15 minutes per sequence for these large ensembles, although we do not, in general, require ensembles one-tenth of this size for analysis. STARLING ensembles were analyzed using the STARLING package (https://github.com/idptools/starling) and using SOURSOP (http://soursop.readthedocs.io/).^76^ Code for generating and analysing ensembles is available on GitHub at https://github.com/holehouse-lab/supportingdata/tree/master/2025/perets_2025.

### All-atom simulations for DDX4^[1-245]^

All-atom Monte Carlo simulations of the N-terminal domain (NTD) of DDX4 were performed using the CAMPARI simulation engine (version 2; http://campari.sourceforge.net/) in conjunction with the ABSINTH implicit solvent model (parameter set: abs_3.2_opls.prm)^77,78^ and the solution ion parameters described by Mao *et al*.^*79*^ Simulations employed the standard CAMPARI movesets and Hamiltonian parameters, as previously reported.^9,80^ Simulations were carried out at 340 K to facilitate robust sampling (as described previously)^9,81^ at 5 mM NaCl in a spherical droplet with a fixed radius of 249 Å to minimize finite-size effects. We note these conditions are somewhat distinct from those examined experimentally, and as such, we focus our analysis of CAMPARI simulations primarily on local backbone behavior over long-range interactions. A total of 20 independent simulations were run, each consisting of 30,000,000 equilibration steps followed by 90,000,000 production steps. During the production phase, conformations were saved every 80,000 steps. Across all simulations, this resulted in a final ensemble comprising 23,773 conformers. CAMPARI simulations were analyzed using SOURSOP.^76^ CAMPARI simulations took ∼1 month per simulation. Self-avoiding random coil simulations (also referred to as the Excluded Volume limit) were performed as described previously.^82^ Input files for simulations, notebooks for simulation analysis, and trajectory information are all available on GitHub at https://github.com/holehouse-lab/supportingdata/tree/master/2025/perets_2025.

### Coarse-grained simulations for DDX4^[1-245]^

Coarse-grained simulations of DDX4 (**Figure S8**) were performed using the LAMMPS simulation and the Mpipi-GG forcefield, a model derived from the Mpipi model.^75,83,84^ Initial configurations for all intrinsically disordered proteins were generated as random walks. Prior to production, up to 1,000 iterations of energy minimization were conducted for each system, continuing until the maximum force dropped below 1 × 10^−8^ (kcal/mol)/Å. Simulations were performed in the NVT ensemble at a NaCl concentration of 150 mM. The temperature was maintained at different values for the coil-to-globule plot using a weakly coupled Langevin thermostat, with adjustments applied every 100 ps. All simulations employed a 20-femtosecond integration timestep. Periodic boundary conditions were used in cubic boxes with a side length of 500 Å. Each simulation was run for 12 milliseconds, with trajectory snapshots saved every 2 µs, generating ensembles of 6000 conformers per temperature. A single simulation per temperature was run, but the extremely long timescale of these simulations, coupled with long gaps between snapshots, means each conformer is effectively independent of the preceding one, as reflected by the extremely smooth coil-to-globule curve. Input files for simulations, notebooks for simulation analysis, and trajectory information are all available on GitHub at https://github.com/holehouse-lab/supportingdata/tree/master/2025/perets_2025.

### Disorder prediction

Disorder prediction (**Figure S1**) was performed using metapredict (V3).^85^

## ACKNOWLEDGEMENTS

The authors acknowledge the late Charles A. Schmuttenmaer (Yale University), in whose laboratory all THz spectroscopy measurements were performed, as well as Dor Ben-Amotz (Purdue University), Rohit V. Pappu (Washington University in St. Louis), Michael K. Rosen (University of Texas Southwestern Medical Center, HHMI), and Shuiqin Zhou (The College of Staten Island, The City University of New York) for insightful feedback during the completion of this work. E.A.P. was supported by the NIH (5T32GM008283-31) and a John C. Tully Chemistry Research Fellowship. J.A.S. acknowledges support from the Onsager Graduate Research Fellowship in Chemistry. J.H.C. was funded by the Science, Technology, and Research Scholars Program at Yale University. A.S.H was supported by a Research Grant from the Human Frontiers Research Program (HFSP) grant RGP0015/2022.

