## Supplementary Information for "Direct Optical Quantification of Chain Collapse, Reduced Dielectric, and Water Release Driving Protein Phase Separation"

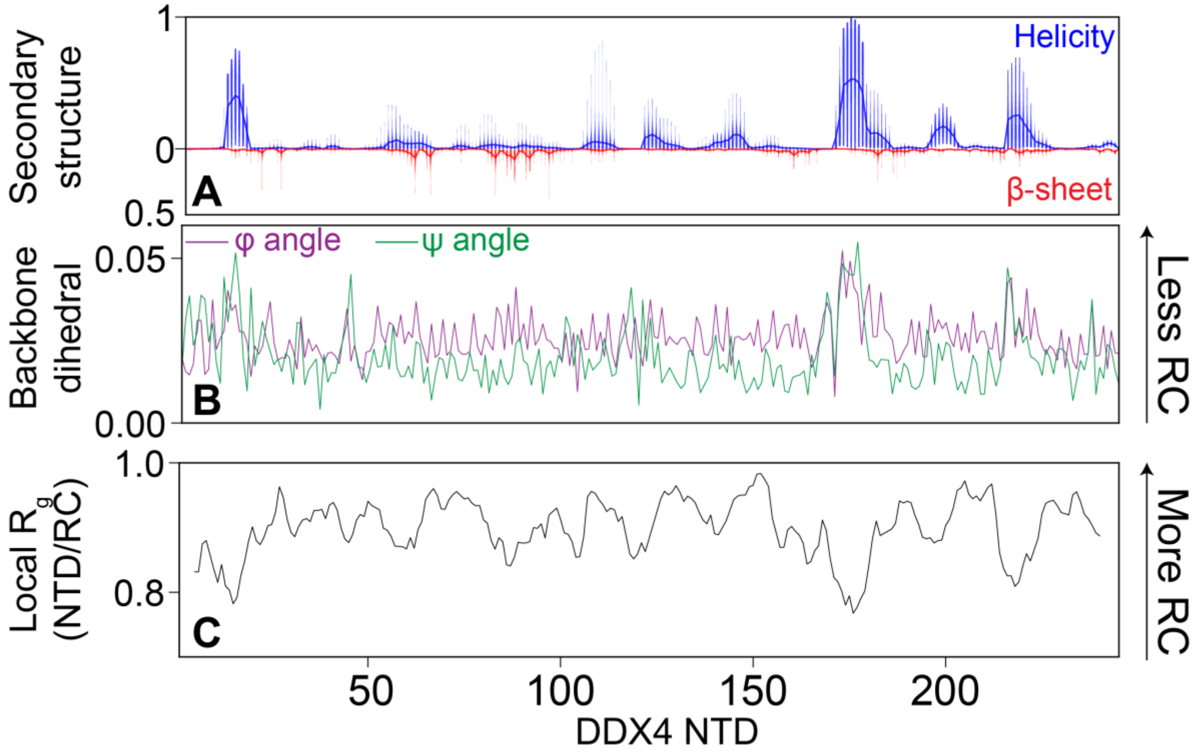


**Figure S1. Assessment of local conformational properties in DDX4[1-245] using all-atom simulations.** (A) For secondary structure assignment, the DSSP algorithm assigns local residues to either helical or ‘extended’ (β-strand/sheet) conformations. (B) For backbone dihedrals, the per-residue value reports on the Hellinger distance between the residue phi (purple) or psi (green) dihedral distributions with the equivalent residue’s distributions from simulations performed in the Self-Avoiding Random Coil limit. Briefly, in self-avoiding random coil simulations, the same all-atom representation is used, but all attractive interactions are set to zero, leading to a conformational ensemble that recapitulates the local and global statistics of a SARC with atomistic detail. The Hellinger distance measures the overall distance between two distributions, where 0 indicates that the two are identical and 1 indicates that they are non-overlapping. The bigger the value, the less random coil (RC) like the residue is. (C) Local conformational behavior is assessed by calculating the radius of gyration for a 10-amino-acid-size “blob” within the chain and dividing by equivalent radius of gyration taken from the SARC simulation. Values closer to 1 are more RC-like, whereas values lower than 1 indicate local intramolecular interactions that lead to a more compact local radius of gyration.

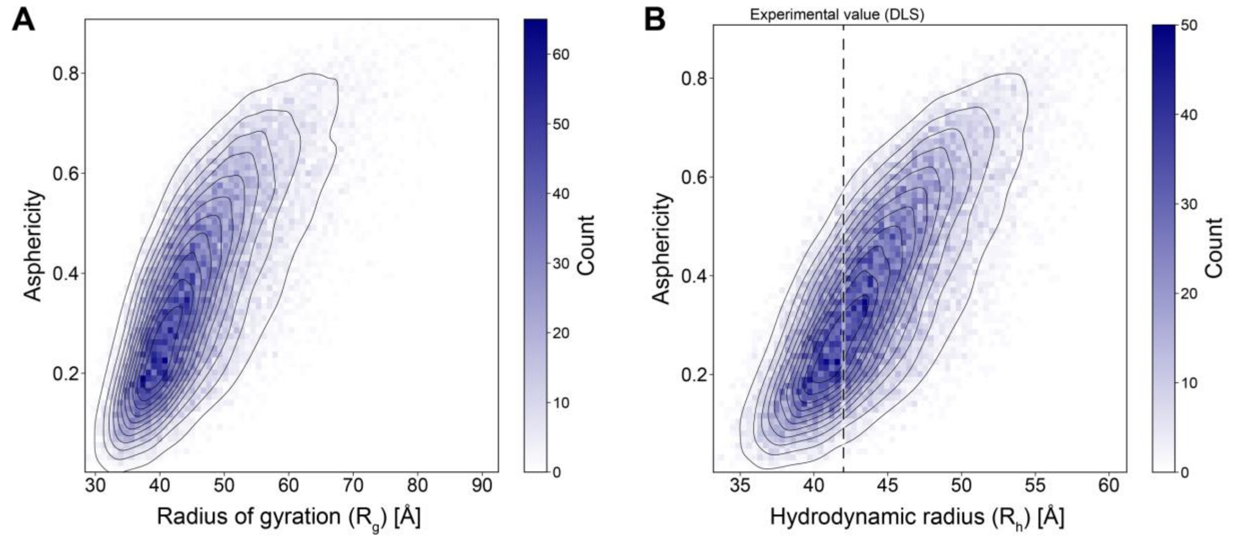


**Figure S2. Assessment of global conformational properties in DDX4[1-245] using all-atom Monte Carlo simulations generated using the ABSINTH implicit solvent model and CAMPARI Simulation Package (ensembles contain 25,000 conformers).** (A) 2D histogram (blue pixels) with Kernel Density Estimates (KDE) (black lines) overlaid for visual clarity describing the 2D distribution of the radius of gyration with asphericity. The average radius of gyration is 45.9 Å, and the broad distribution is consistent with a well-sampled highly disordered protein. (B) 2D histogram with KDE lines overlaid for visual clarity describing the 2D distribution of the hydrodynamic radius with asphericity. The average hydrodynamic radius is 44.03 Å, in reasonable agreement with the experimental value of 42 Å obtained from DLS. This broad distribution is consistent with a well-sampled, highly disordered protein.

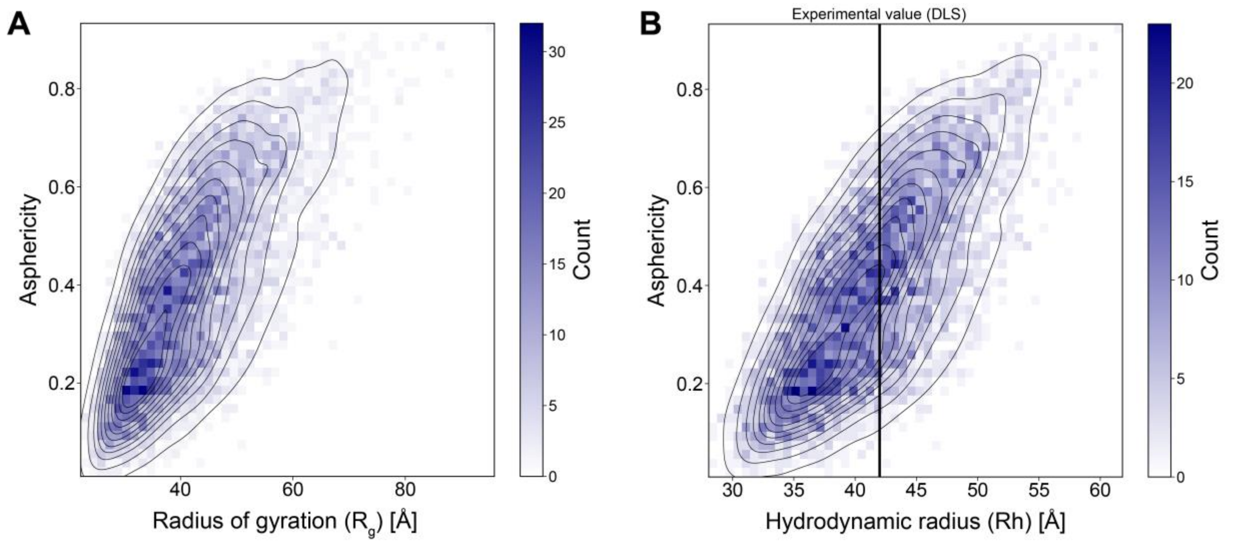


**Figure S3. Assessment of global conformational properties in DDX4[1-245] using STARLING-derived ensembles (ensembles contain 6000 conformers).** (A) 2D histogram (blue pixels) with Kernel Density Estimates (KDE) (black lines) overlaid for visual clarity describing the 2D distribution of the radius of gyration with asphericity. The average radius of gyration is 41.7 Å, and the broad distribution is consistent with a well-sampled highly disordered protein. (B) 2D histogram with KDE lines overlaid for visual clarity describing the 2D distribution of the hydrodynamic radius with asphericity. The average hydrodynamic radius is 41.4 Å, in good agreement with the experimental value of 42 Å obtained from DLS. This broad distribution is consistent with a well-sampled, highly disordered protein.

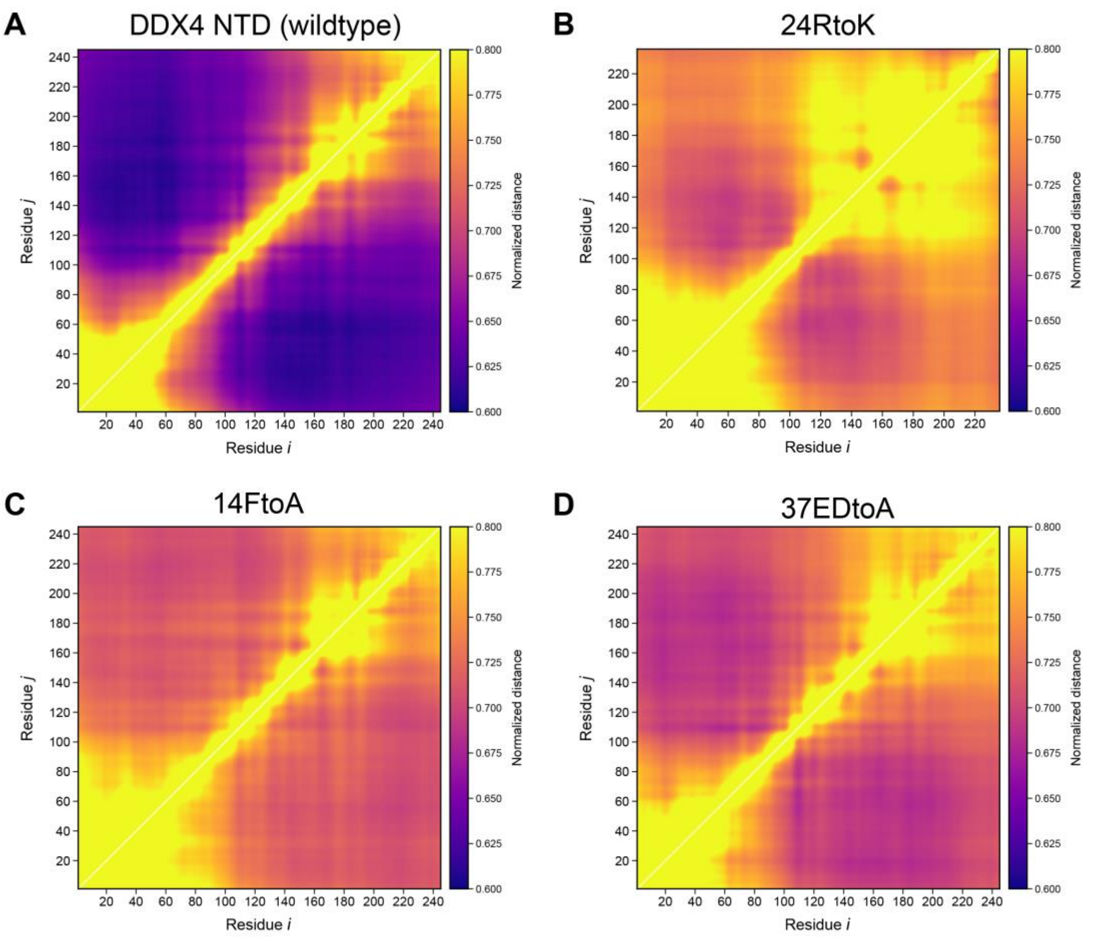


**Figure S4. Intramolecular scaling maps for wildtype and DDX4 NTD variants.** Scaling maps are calculated as the ratio of the distance seen in the specific variant, normalized by the distance seen for a serine-proline dipeptide repeat as a null model for an expanded, uniformly interacting IDR. (A) DDX4$^{[1\text{-}245]}$ scaling map reveals long-range interactions that deviate from an expanded chain (blue regions) across the sequence; notably, residues between 80 and 140 contribute extensive long-range interactions. (B, C, D) A marked and overall reduction in long-range interactions is observed in the 24RtoK, 14FtoA, and 37DEtoA variants, as indicated by an increase in the normalized distances. Note the 24RtoK variant uses the DDX4$^{[1\text{-}236]}$ construct to match previously reported experimental work.

Using intrachain scaling behavior, an estimate of the apparent Flory scaling exponent ($\nu^{app}$) for CAMPARI and STARLING derived ensembles is 0.51 and 0.52, respectively.[1] While interpreting an apparent scaling exponent has many caveats, both values are consistent with chains in marginally good solvents, close to scaling behavior expected for a polymer in a theta solvent ($\nu = 0.5$). These results are consistent with a model in which DDX4$^{[1\text{-}245]}$ is poised for environmental control of solvent quality. Indeed, the original characterization of the DDX4 NTD focused on its environmental sensitivity with respect to phase separation.[2] When conditions change to those that promote phase separation (decreased salt or temperature), monomer collapse is driven by the reduction in solvent quality. These considerations lead us to consider a model of DDX4 NTD phase separation where intramolecular and then intermolecular interactions are favored, first leading to monomer collapse and ultimately to phase-separated assemblies (**Figure 6**, main text).

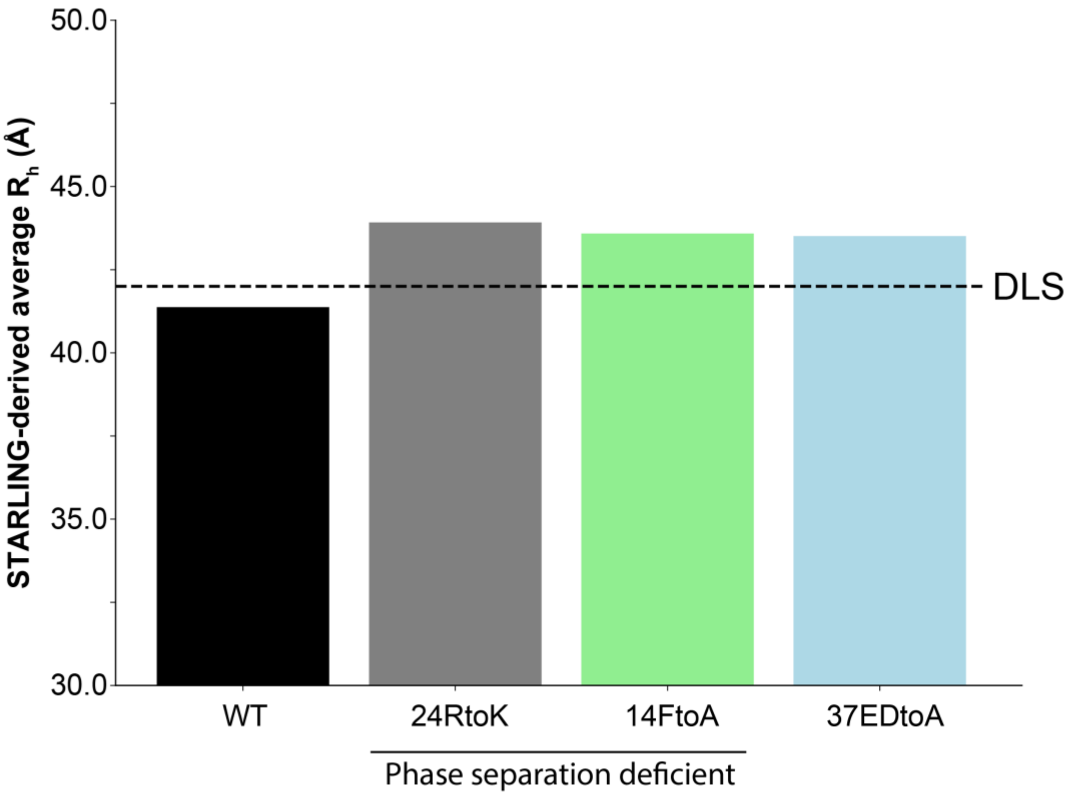


**Figure S5. Predicted hydrodynamic radii from DDX4 NTD variants.** In all cases, the loss of long-range intramolecular interactions (see **Figure S4**) is accompanied by an increase in overall global dimensions, as reported here by an increase in the hydrodynamic radius ($R_h$).

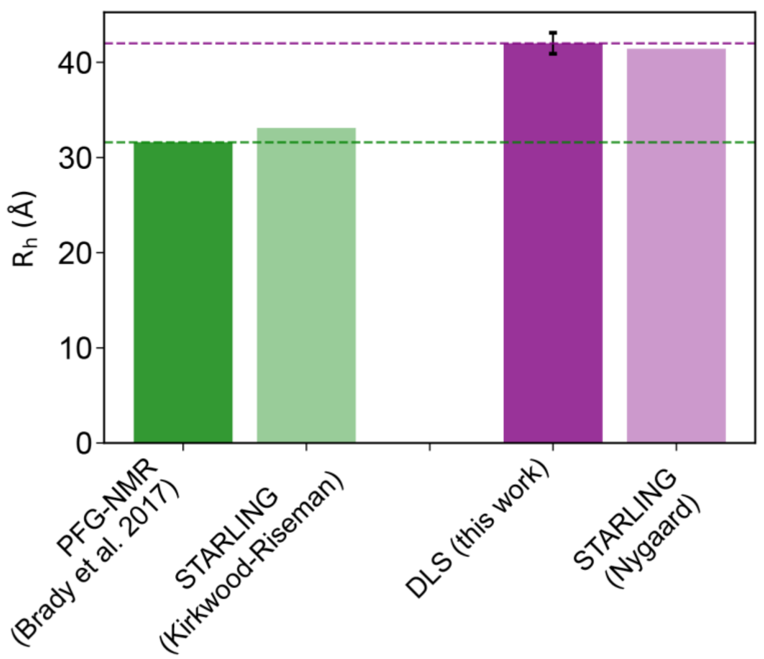


**Figure S6. Comparison of experimental and STARLING-ensemble derived hydrodynamic radii ($R_h$) values obtained from this and prior work, using distinct methodologies.** Prior work by Brady et al. using Pulse Field Gradient Nuclear Magnetic Resonance (PFG-NMR) spectroscopy reported a hydrodynamic radius of 3.16 nm, in extremely good agreement with the $R_h$ derived from STARLING ensembles using the Kirkwood-Riseman approximation.[3] The Kirkwood-Riseman approximation has previously been suggested to offer better agreement with PFG-NMR-derived hydrodynamic radii, in agreement with our findings here.[4] In contrast, the $R_h$ obtained from DLS here is, in principle, larger, at 4.2 nm; yet, for the same STARLING ensemble, an $R_h$ of 4.14 nm is recovered using the approximation of Nygaard et al.,[5] which offers good agreement with our DLS data.

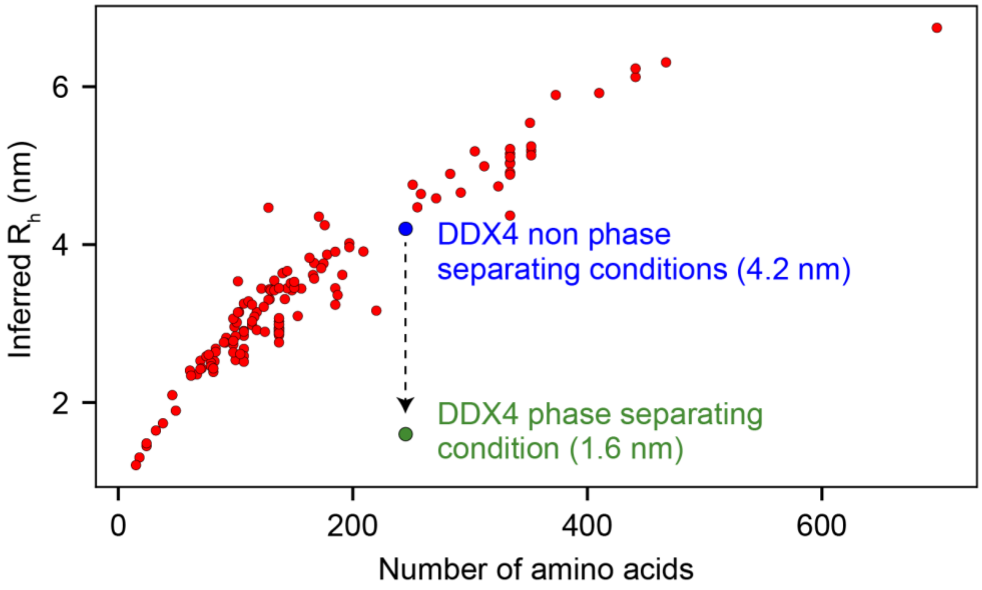


**Figure S7. Comparison of inferred *$R_h$* for 144 proteins** (values originally reported in Alston *et al.*).[1] $R_h$ is calculated by taking extant $R_g$ values obtained from SAXS data and converting $R_h$ using the approximation of Nygaard *et al.*[5] Superimposed are our experimentally determined $R_h$ values for DDX4$^{[1\text{-}245]}$ under non-phase separating conditions (blue) and under phase separating conditions (green) (Figure 3, main text).

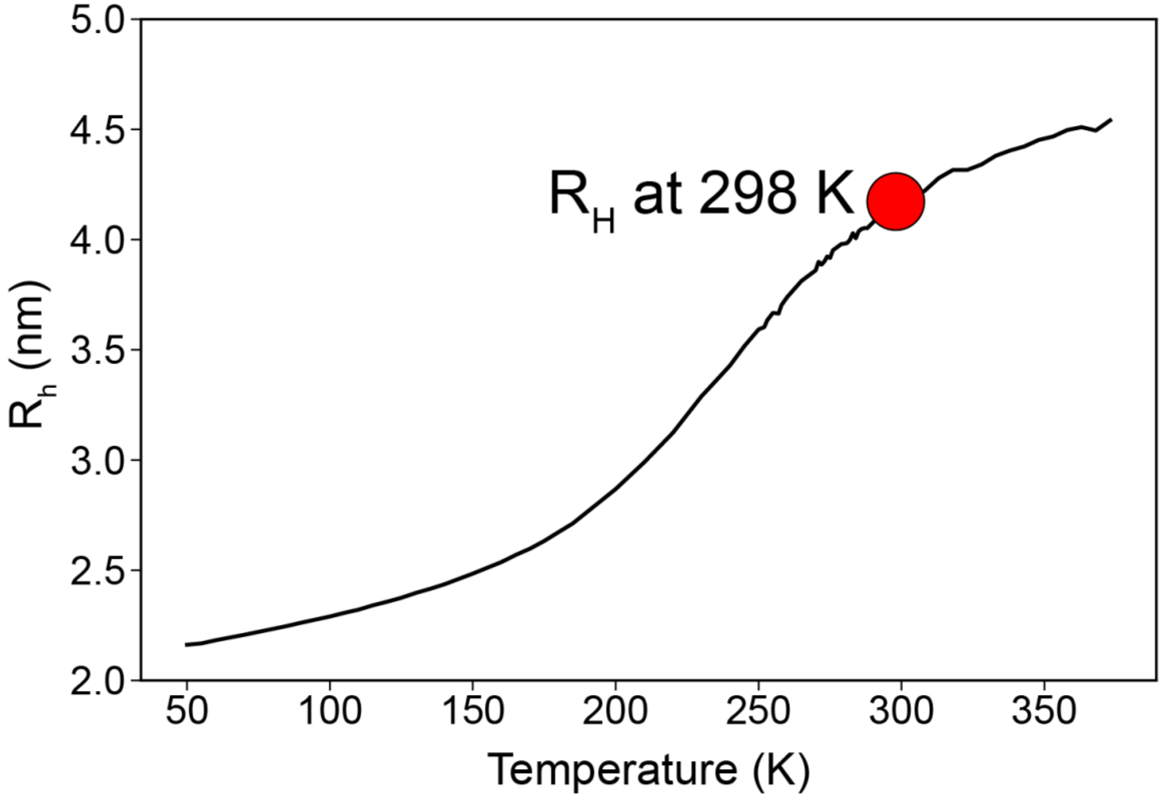


**Figure S8. Theoretical coil-to-globule curve for DDX4$^{[1-245]}$ obtained from coarse-grained molecular dynamics simulations performed using the LAMMPS simulation engine and the Mpipi-GG forcefield.** The curve here reflects a temperature-dependence in the context of the Mpipi-GG model,[6-8] but temperature should be treated here as a parameter scaling the strength of intramolecular interactions as opposed to *bona fide* temperature.

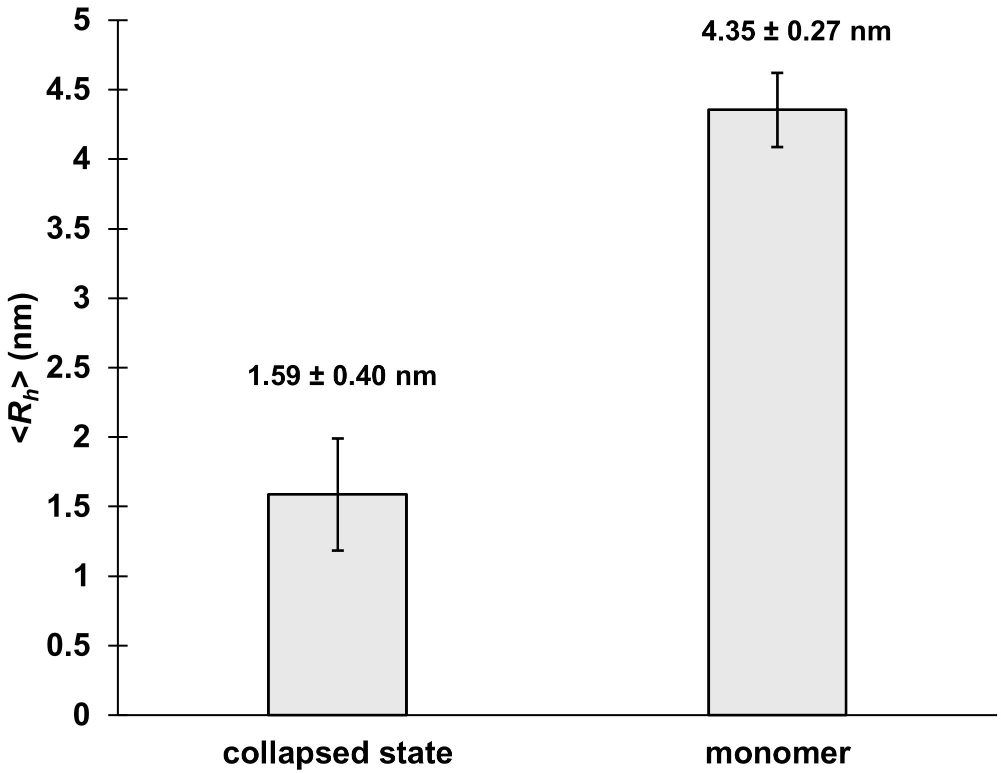


**Figure S9. Dynamic light scattering reveals a reproducible collapsed state of DDX4[1-245] in phase separation.** Average $R_h$ from $N$ = 11 experiments measured at 200 mM NaCl and 50 µM DDX4[1-245] at 10°C. Error bars indicate one standard deviation.

**Table S1.** Dielectric permittivity fit parameters for samples in $H_2O$.

| | $\varepsilon_s$[a] | $\varepsilon_\infty$ | $\Delta\varepsilon_1$ | $\Delta\varepsilon_2$ | $\tau_2$ (ps) | $A_S$ |
|---|---|---|---|---|---|---|
| *Water* | 84 ± 4 | 2.53 ± 0.08 | 77 ± 5 | 2.2 ± 0.5 | 0.3 ± 0.1 | 39 ± 7 |
| *Buffer* | 84 ± 4 | 2.5 ± 0.1 | 78 ± 5 | 2.1 ± 0.5 | 0.27 ± 0.08 | 36 ± 9 |
| *Dilute* | 78 ± 8 | 2.4 ± 0.2 | 71 ± 9 | 2.6 ± 0.8 | 0.3 ± 0.1 | 45 ± 7 |
| *Condensed* | 67 ± 6 | 2.58 ± 0.08 | 61 ± 7 | 2.5 ± 0.6 | 0.3 ± 0.1 | 39 ± 8 |

[a] The static permittivity ($\varepsilon_s$) is a calculated parameter.

**Table S2.** Dielectric permittivity fit parameters for samples in $D_2O$.

| | $\varepsilon_s$[a] | $\varepsilon_\infty$ | $\Delta\varepsilon_1$ | $\Delta\varepsilon_2$ | $\tau_2$ (ps) | $A_S$ |
|---|---|---|---|---|---|---|
| *Buffer* | 71 ± 4 | 2.6 ± 0.4 | 65 ± 4 | 2.3 ± 0.2 | 0.26 ± 0.03 | 29 ± 11 |
| *Dilute* | 69 ± 4 | 2.4 ± 0.5 | 63 ± 4 | 2.0 ± 0.3 | 0.27 ± 0.04 | 38 ± 16 |
| *Condensed* | 49 ± 2 | 2.0 ± 0.4 | 44 ± 3 | 2.1 ± 0.7 | 0.2 ± 0.1 | 41 ± 8 |

[a] The static permittivity ($\varepsilon_s$) is a calculated parameter.
